# Cryogenic light microscopy of vitrified samples with Ångstrom precision

**DOI:** 10.1101/2025.05.27.656160

**Authors:** Hisham Mazal, Franz-Ferdinand Wieser, Daniel Bollschweiler, Vahid Sandoghdar

## Abstract

High-resolution studies in structural biology are commonly based on diffraction methods and on electron microscopy. However, these approaches are limited by the difficulty in crystallization of biomolecules or by a low contrast that makes high-resolution measurements very challenging in crowded samples such as a cell membrane. The exquisite labeling specificity of fluorescence microscopy gets around these issues. Indeed, several recent reports have reached resolutions down to the Ångstrom level in super-resolution microscopy, but to date, these works used fixed samples. To establish light microscopy as a workhorse in structural biology, two main requirements must be fulfilled: near-native sample preservation and near-atomic optical resolution. Here, we demonstrate a technique that satisfies these key criteria with particular promise for conformational studies on membrane proteins and their complexes. To prepare cell membranes in their near-native state, we adapt established protocols from cryogenic electron microscopy (Cryo-EM) for shock-freezing and transfer of samples. We developed a high-vacuum cryogenic shuttle system that allows us to transfer vitrified samples in and out of a liquid-helium cryostat that houses a super-resolution fluorescence microscope. Sample temperatures below 10 K help dissipate the heat from laser illumination, thus maintaining intact vitreous ice. We utilize the photoblinking of organic dye molecules attached to well-defined positions of a protein to localize one label fluorophore at a time. We present various characterization studies of the vitreous ice, photoblinking behavior, and the effects of the laser intensity. Moreover, we benchmark our method by demonstrating Ångstrom precision in resolving the full assembled configuration of the heptameric membrane protein alpha-hemolysin (αHL) in a synthetic lipid membrane as a model system. Additionally, we report on the technique’s capability to resolve membrane proteins in their native cellular membrane environment. Our method, which we term single-particle cryogenic light microscopy (spCryo-LM), enables structural studies of membrane protein tertiary and quaternary conformations without the need for chemical fixation or protein isolation. The approach can also integrate other super-resolution or spectroscopic techniques with particular promise in correlative microscopy with images from Cryo-EM and related techniques.

## Introduction

Imaging is desirable in nearly all aspects of cell biology, but arguably one of the most challenging tasks of high-resolution microscopy is to shed light on the behavior of membrane proteins. Indeed, intensive efforts have been dedicated to the investigations of protein structure and function (1–4), but the intricate details of proteins and their complexes in the cell remain difficult to access. Advances in sample preparation (*5*) and cutting-edge hardware have sparked a revolution in cryo-electron microscopy (Cryo-EM), reaching atomic resolution in isolated protein macromolecules (*6–9*). However, a complete insight into protein function can only be achieved in the native ultrastructural cellular context to ensure proper consideration of all biochemical and physical interactions. This ambitious goal is currently being pursued by cryogenic electron tomography (Cryo-ET) as a new variant of Cryo-EM (*10*), where protein structures and conformations are studied within cells and tissues preserved in their near-native states (*11–13*). A major drawback of Cryo-EM methods is that they necessitate a low-dose electron beam to avoid radiation damage, resulting in low signal-to-noise ratios (SNR). Thousands of individual particle images are usually averaged and classified in order to reach atomic resolution (*10, 14*). Consequently, the technique still faces significant challenges in identifying target molecules in their native state (*10, 15, 16*). For example, quantitative insights into the structure of integral membrane proteins (*17–19*), which account for 20-30% of the total proteome (*20, 21*), and their organization within the cell membrane, remain scarce (*22–24*).

Fluorescence microscopy provides molecular specificity and exceptional sensitivity down to the single molecule level, making it an established workhorse for studying cellular and sub-cellular structures. The remarkable recent progress in super-resolution microscopy has pushed the three-dimensional (3D) spatial resolution to 15-20 nm in routine studies (*25*), and state of the art techniques such as minimal photon flux (MINFLUX) (*26*) and DNA-PAINT (*27*) have even reported precision on the sub-nanometer scale. Because this level of spatial resolution requires exceptional mechanical stability to counter unwanted effects caused by thermal molecular jitter, instrumental vibrations and drifts, samples have been chemically fixated for room temperature (RT) investigations. The fact that chemical fixation can introduce artifacts and distort the system under study, however, makes this approach suboptimal (*28–31*).

To date, shock-freezing is considered to be the method that best preserves cellular and molecular structures, providing near-native state preservation of biological samples with minimal disruption (*32, 33*). Having being invented in the context of Cryo-EM, this technique has also been used for cryogenic fluorescence microscopy, but mainly to help identify target proteins prior to measurements in an electron microscope (*34*). The majority of such setups operate at the liquid nitrogen (LN) temperature and atmospheric pressure (*35–37*), suffering from mechanical and thermal instabilities as well as condensation and devitrification (*35*); see recent review for details (*34, 38*). The central challenges in these works are: 1) laser illumination is restricted to low intensities in order to prevent sample devitrification (*39*); 2) photophysical properties of fluorophores at low temperatures remain poorly understood (*36, 37, 40*); 3) constrained conformational changes of fluorescent proteins under cryogenic conditions reduce the prospects of photoactivation for localization microscopy (*34*); 4) applicability of special chemicals that enhance photoblinking is less effective due to hindered diffusion at low temperatures. As a result, optical resolution has generally remained in the range of 10-100 nm in these methods. A recent work pushed cryogenic light microscopy to a higher localization precision in vitrified samples prepared on thick sapphire discs at liquid helium temperature (LHe) (*31*). However, this choice of substrate is not favorable for direct correlative Cryo-EM studies.

In this work, we report on an approach for resolving protein conformation and assembly with Ångstrom precision in their native state. By using commercial transmission electron microscopy (TEM) grids as sample carrier, we devised a method that is readily compatible with Cryo-EM and correlative imaging. To achieve this, we engineered a high-vacuum cryogenic shuttle system that allows us to transfer shock-frozen vitrified samples in and out of a cryogenic optical microscope with minimal condensation and devitrification. We benchmark our method on a model system consisting of the heptameric membrane protein, alpha-hemolysin (αHL), that is reconstructed in small unilamellar vesicles, which are in turn used to create a synthetic lipid bilayer (SLB). We also report on the ability of our pipeline to interrogate transmembrane proteins in their native cellular membrane. The work flow developed here can also be used for investigations of other cell biological samples with a variety of optical techniques such as Raman spectroscopy (*41*).

## Results

### Cryogenic light microscopy of vitrified samples

Several years ago, we introduced cryogenic localization microscopy in three dimensions (COLD), in which Ångstrom resolution was achieved by exploiting the stochastic photoblinking of single fluorescent labels on isolated protein molecules embedded in a polymer (*42, 43*). In the current work, we modify the experimental process to allow for measurements on biological samples that are preserved in their near-native state via shock freezing. Given that the method allows one to visualize the organization of single proteins, we also refer to it as single-particle cryogenic light microscopy (spCryo-LM) in analogy to single-particle cryogenic electron microscopy (spCryo-EM).

One of the common techniques for vitrification entails rapid immersion of a TEM grid, which supports a sample covered by a thin layer of water, into a cryogenic liquid (*5*). This process, called plunge freezing, achieves sufficiently high cooling rates for a sample thickness up to about 1 µm (*44*) and enables the preservation of macromolecules, protein crystals, viruses, cells, and even thin bacteria in their near-native states (*45, 46*). Subsequent to plunge freezing, the vitrified sample must be maintained below the critical temperature of 130 K to prevent ice re-crystallization (*44*). Additionally, exposure to air must be minimized to reduce contamination arising from condensation on the cold surface. To fulfil these conditions, we adapted a design strategy from Cryo-EM applications for use in our setup (*47*), which allow us to transfer vitrified specimens between different apparatuses at high vacuum and LN temperature to minimize condensations and devitrification Figure 1 sketches the principal workflow of shock freezing the sample and its transfer from a preparation chamber into the cryogenic optical microscope. We first plunge freeze the sample in liquid ethane and then store it in a dedicated TEM grid box (Fig. 1a, positions 1-3). Next, the grid box is docked onto a sample-handling platform in an insulated glass vessel filled with LN inside a preparation chamber (Fig. 1b, positions 3-6). The platform is held by two polyether ether ketone (PEEK) isolation brackets connected to the top flange (Fig. 1b, and Supplementary Fig. 2). The glass vessel is attached to a vacuum-tight vertical manipulator (Fig. 1b, position 7) that allows us to move it up and down during the transfer process. Figure 1b (i) shows a close-up view of the sample-handling platform. A groove helps hold the TEM grid box (position 3), a cold stage with a dovetailed structure (position 4) supports a sample cartridge (position 5, Fig. 1b(ii, iii)). An anti-contaminator plate (position 6) can be slid over the sample to protect it during transfer (Fig. 1b (i)). The TEM grid is picked up *in situ*, i.e. in LN, from the grid box and is mounted onto the sample cartridge and secured with a thin-profile clamp, using a small screw (Fig. 1b (iii), and Supplementary Fig. 1-3). To ensure good contact between the sample cartridge and the cold stage, we used two pairs of repelling magnets, press-fitted to the bottom of both the cartridge and the cold stage (Supplementary Fig. 1). A sensor is attached to the complementary holder close to the sample cartridge to monitor the chamber temperature (Supplementary Fig. 2).

**Fig. 1.**
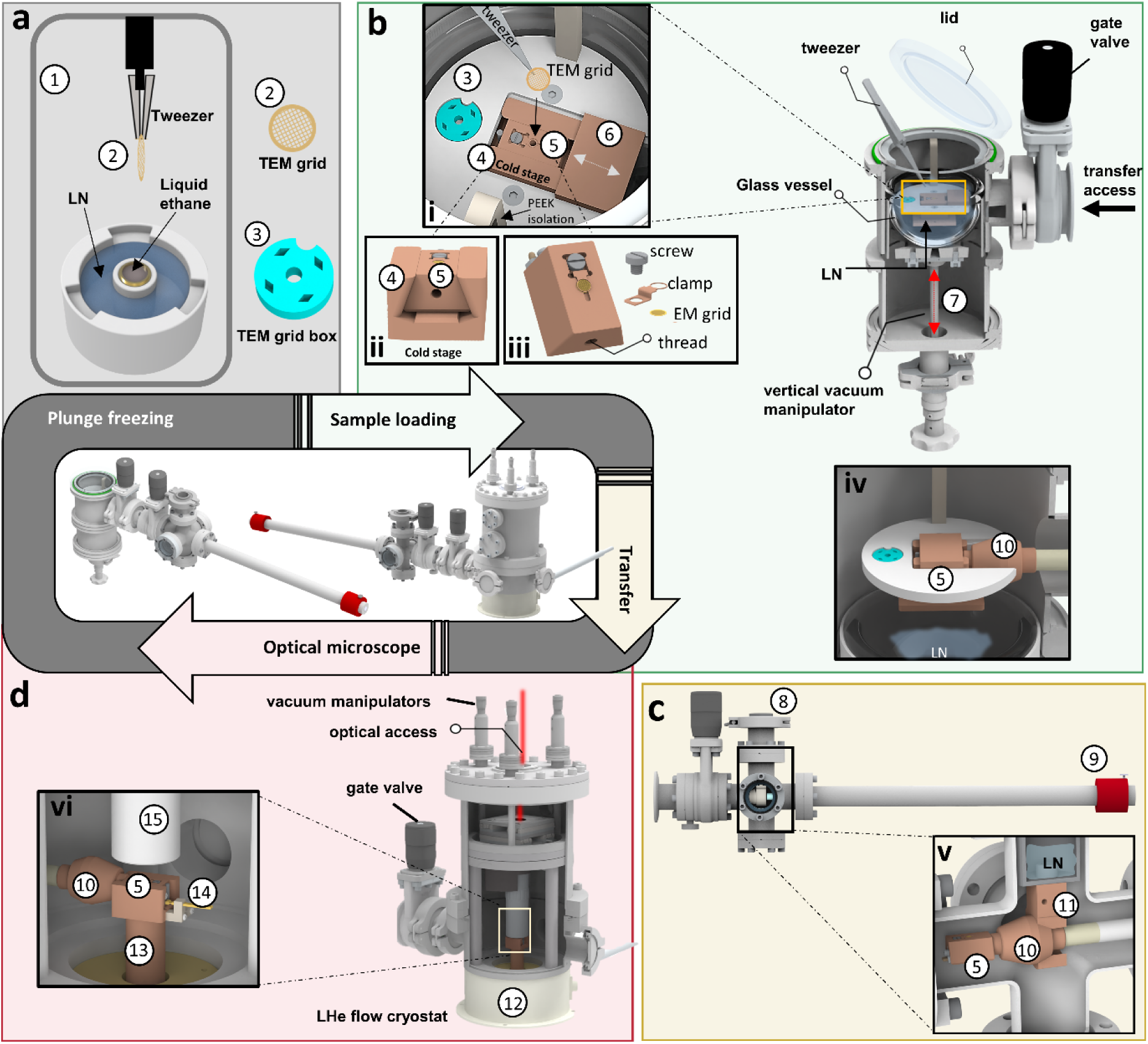
Cryogenic sample preparation and transfer. **a**, Plunge freezing is achieved in a dedicated commercial instrument containing a cryogenic liquid (1). The sample on a TEM grid (2) is then placed and stored in a TEM grid box filled with LN (3). **b**, A sample loading chamber uses a glass vessel to surround a cold stage (4) that holds a sample cartridge (5) with LN. The vessel can be lowered by a vacuum-tight manipulator (7). A lid can close the chamber for application of vacuum. A gate valve allows access from the side for the sample transfer. After the grid box (3) is transferred, the TEM grid is loaded and clamped onto the sample cartridge (see inset i-iii). A sliding plate acts as an anti-contaminator cover (6). **c**, A vacuum-tight transfer shuttle can be flanged to the loading chamber and the cryostat via a gate valve. The essential components include a small LN dewar (8) and a vacuum-compatible manipulator (9). Inset (iv) shows the connection to the preparation chamber where the transfer shuttle is loading the sample cartridge after lowering the LN vessel. Inset of (v) shows a cross-section of the LN dewar. The manipulator rod fits through a c-shaped cold stage (11), and a conical taper receives the head (10), providing a perfect fit for efficient thermal conduction to the sample cartridge (5). **d**, The transfer shuttle is connected to the cryostat which houses the optical microscope (12). Inset vi shows the connection of the cartridge carried by the transfer shuttle with a dedicated cold finger (13) in the cryostat. The microscope is based on a Janis-500 cryostat (12) with a series of modifications to accommodate a 3D piezo scanner and high-NA microscope objective (100 x, 0.9 NA, 2mm working distance) in vacuum (15, inset vi) (*42*). The cold finger (13) was fitted with a dove tail to receive the sample cartridge, as well as a temperature sensor reader (14).

Upon loading the TEM-grid, we slide the anti-contaminator plate (Fig. 1b, position 6) onto the top of the sample cartridge to block any condensations or contaminations. Next, we close the lid of the preparation chamber, and lower the glass vessel containing LN to prepare the transfer of the cartridge to the side (Fig. 1b-c). The chamber is then evacuated to 10^0^-10^-1^ mbar, as some of LN remains in the glass vessel and does not fully vaporize within the timeframe of the transfer process (see Supplementary Method 1, Supplementary Fig. 2, Movie 1, and Supplementary protocol). To suppress water vapor in the preparation chamber, we de-gas the preparation chamber with dry nitrogen throughout the entire process up until the evacuation step, during which the sample cartridge remains fully immersed in liquid nitrogen. A manipulator from a high-vacuum shuttle (1×10^-6^ mbar) operating at LN temperature is used to transfer the cartridge from the loading chamber (Fig. 1c) to a cryostat that houses the optical microscope pre-cooled at LHe temperature (Fig. 1d). Figure 1c shows the full transfer shuttle. The essential components include a small LN dewar (position 8) and a linear vacuum-compatible manipulator (position 9). The manipulator rod contains a large-mass conical head (position 10) made from oxygen-free copper, which holds a screw to load the sample cartridge through a threaded hole on its side (Fig. 1b (iii) and Supplementary Fig. 2).

After loading the sample cartridge (position 5) from the preparation chamber into the transfer shuttle under low vacuum, the cartridge is brought to contact with the LN temperature cold stage (position 11), via the tool head. This enables the sample to maintain its temperature below the devitrification point (Supplementary Fig. 2). Next, the gate valve of the preparation chamber is closed to allow the transfer shuttle to reach high vacuum. Then, the gate valve of the transfer shuttle is closed, the transfer shuttle is disconnected from the preparation chamber, and it is connected to the cryostat through a KF50 flange (Fig. 1d). Then, we transfer the cartridge to the cold stage inside the optical microscope, which is kept under high vacuum and cooled to LHe temperature (Fig. 1d, position 13). A comprehensive workflow, characterization and description is provided in the Supplementary Method 1, Supplementary Figs. 1-4, Movie 1, and Supplementary protocol. The procedure can also be executed in the reverse order for grid extraction and subsequent analysis and correlative imaging.

To obtain sufficiently large numbers of photons during each blinking on-time, it is desirable to excite a molecule at a high rate, requiring laser intensities in the range of 0.6 - 1 kW/cm^2^. This level of light intensity has been reported to cause severe sample devitrification at 77 K, especially in the case of carbon mesh TEM grids (*36, 39*). We, thus, screened several grid materials, including carbon and gold mesh structures with various shapes at 8 K. We found that carbon meshes are unstable, buckle and tear at high laser intensities ∼ 0.3-0.6 kW/cm^2^ (Supplementary Fig. 5 and Movie 2). As a result, we opted to work with UltrAufoil TEM grids R2/2 200 mesh (UF), which have been shown to offer superior mechanical stability (*48*) and to handle high optical intensities (∼ 0.6 kW/cm^2^) (*39*), particularly when using light in the red spectral range (*49*), thereby maintaining stable vitreous ice condition (Supplementary Fig. 5).

To investigate the quality of the entire vitrification procedure and the effect of laser illumination, we examined the vitrified TEM grids using Cryo-EM. To validate proper blotting vitrification with a visual check, we added ∼ 1 nM ATTO647N dye molecules to pure aqueous samples (Supplementary Method 2, and Supplementary Fig. 6). We exposed different grids to varying laser intensities (0, 0.1, 0.3,0.6, and 1 kW/cm^2^) in the cryostat. Each field of view was illuminated for approximately 15-20 minutes (Supplementary Fig. 6 and Supplementary Method 2). The samples were subsequently retracted from the cryostat and examined using Cryo-EM imaging. Micrographs with a total electron dose of 60 e^-^/ Å² at an equivalent magnification of 0.85 Å/px were recorded. The occurrence and intensity of crystalline ice diffraction rings is generally be used to assess the presence of unwanted crystalline ice, with characteristic bands between 6 and 30 Å (*50, 51*). The calculated power spectra of the vitreous ice exposures in holes, i.e., the modulus of their Fourier transforms to the second power, is used as a quantitative measure for the presence of devitrified ice. We found that only the 1 kW/cm^2^ pre-exposure unambiguously exhibited notable devitrification in the exposed region (Supplementary Fig. 6, Methods, and Supplementary Method 2).

### Photophysics characterization

Having validated that our sample remains vitrified during laser illumination, we sought to characterize photoblinking in vitreous ice since photophysics of molecules is known to be highly sensitive to their material environment (*52*). Here, we recorded the fluorescence from a vitrified solution of ATTO647N in water on UF TEM grids. As depicted in Fig. 2a, molecules were spread sparsely on the TEM grid and show high photon counts. Moreover, Fig. 2b,c shows that we observe fast spontaneous blinking with an average off-on time ratio of ∼ 30, as a result of longer off-times than the on-times (Movies 3-4). This feature enables resolving of multiple fluorophores below the diffraction limit (*42, 53*). Some previous studies reported weak blinking behavior at LN temperature, both under atmospheric pressure and in high vacuum (*37, 54*) while other work at high vacuum and LHe temperature (*31*) report strong photoblinking in line with what was observed in our work.

**Fig. 2.**
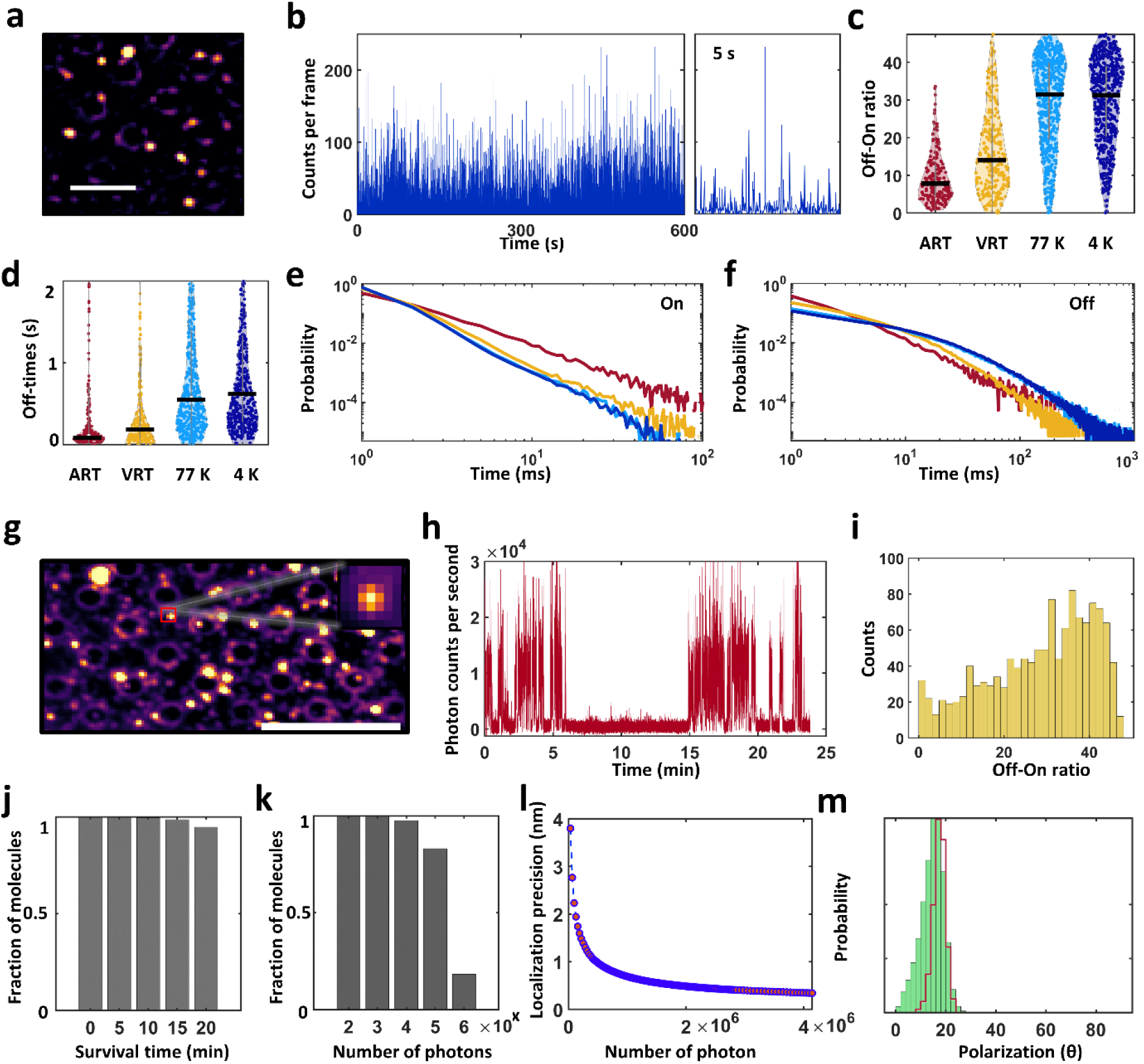
Photophysics characterization of fluorescent molecules in vitreous ice. **a**, Fluorescence image of a vitrified aqueous solution of ATTO647N recorded on a UF TEM grid. Scale bar is 5 µm. **b**, Exemplary intensity time traces of a single dye molecule show large off-times and short on-times suitable. Right panel shows a close-up of a 5 second interval. **c**, Off-on ratios obtained from a large number of molecules at different conditions. ART: ambient room temperature; VRT: vacuum room temperature; 77K: vacuum, LN temperature; 4K: vacuum, LHe temperature. The black horizontal lines indicate median values. Black lines indicate median values. **d**, Same as (d) but for off-time distributions. **e-f**, Probability distributions of the on and off times presented on log-log plots. Color code is same to that in (c,d). **g**, smURFP protein imaged in vitreous ice. Scale bar is 10 µm. **h**, Intensity time trace shows exceptionally long off-times. **i**, Off-on ratio histogram as obtained from many molecules. **j-l**, Survival time histogram shows that > 90% of the molecules survived for at least 10 min, allowing the collection of > 1 x10^6^ photons and, thus, a localization precision down to the Ångstrom level. Localization precision better than 1 nm can be achieved if one uses > 3000 localizations events. **m**, Polarization histogram of a single molecule indicating a stable and narrow distribution in vitreous ice, suitable for distinguishing individual molecules based on their orientations. The histogram becomes narrower (red line) when filtering the data to include only frames with a high photon count.

Next, we discuss the effect of vacuum and temperature. First, we imaged the ATTO647N sample at RT and ambient atmospheric pressure (ART). Then the same sample was imaged at RT but under high vacuum (VRT), followed by LN and high vacuum, and finally by LHe and high vacuum. As depicted in Fig. 2c, we found that a moderate vacuum (∼10⁻⁴ mbar) significantly enhances the off-on ratio as compared to atmospheric pressure by about a factor of two. Moreover, we observed that decreasing the temperature from RT to LN or further down to 8 K enhanced photoblinking by roughly another factor of two relative to ART. By analyzing the off-dwell and on-dwell times (Supplementary Fig. 7), we found that this enhancement results from both an increase in off-times and a decrease in on-times. As depicted in Fig. 2d, the median off-time increases as a function of pressure by a factor of 4, as a function of LN temperature by an additional factor of 3.2, and slightly improved by an additional factor of 1.1 at 8 K. We note that our estimate of the on-times is limited to our frame rate of ∼ 14 ms. We also found similar effects in other far-red organic dyes such as Cy5, iFluor 647 and Alexa Fluor 647 (Supplementary Fig. 7). The on-time distribution (Fig. 2e) indicates a power-law behavior with an increasing power exponent as the temperature decreases. The off-time distribution (Fig. 2f) also follows a power-law behavior at RT, but it shifts to a biexponential behavior as vacuum is applied and the temperature is reduced.

In addition to organic molecules, we also tested the photophysics of far-red fluorescent proteins, such as small Ultra Red Fluorescent Protein (smURFP)(*55*) and observed a similar behavior (Fig. 2g-I, Movie 5). We emphasize that this blinking behavior is completely spontaneous, i.e., without the use of any special buffer conditions as required in techniques such as stochastic optical reconstruction microscopy (STORM)(*56*). In summary, while not all photoblinking effects at cryogenic temperatures are yet fully understood, our work shows that combination of vacuum and low temperature provides favorable conditions for the use of spontaneous photoblinking in single-molecule localization microscopy at cryogenic temperature.

Another advantage of cryogenic temperatures, besides superior sample preservation and favorable photoblinking, stems from the fact that fluorophores remain stable against photobleaching for extended periods of time (Fig. 2j). This allows one to detect several orders of magnitude more photons from a single molecule than at ambient conditions (Fig. 2k). It is, indeed, this feature that is responsible for localization precisions in the range of a few Ångstroms (Fig. 2l).

### Super-resolution imaging in vitrified samples

In order to resolve multiple fluorophores below the diffraction limit, one needs to isolate and localize each fluorophore separately. In our approach, we exploit the fixed dipole orientations of fluorescent molecules at low temperature to distinguish them (*43, 53*) based on their stochastic photoblinking which allow us to sperate them over time. To do this, the dipole orientation of a molecule is projected onto the angular interval between [0°-90°] after its fluorescence signal traverses a polarizing beam splitter, and its orientation is determined in a ratiometric manner from the intensities of the resulting point-spread functions (PSF) on two cameras. (*53*). Figure 2m plots an exemplary histogram of the polarization signal recorded from a single fluorophore.

### DNA nanoruler

As a first demonstration, we imaged DNA origami nanorulers that contained two ATTO647N dye molecules separated by 30 nm (Fig. 3a). We loaded 3 μl of this sample onto a UF TEM grid, followed by plunge-freezing. Fluorophores are free to rotate before shock freezing, resulting in random 2D projections (Supplementary Fig. 8). The vitrified sample is transferred and imaged using the cryogenic optical microscope (Movie 6). Figure 3b showcases a polarization signal with two polarization components at ∼ 20° and 75°, and Fig. 3c shows their corresponding PSFs. Co-localization of the two components (see method section) allows us to generate a super-resolved image that measures the fluorophore positions with Ångstrom precision (Fig. 3d). Figure 3e depicts the distance histogram obtained from 55 different projections (N = 55). The histogram shows a main peak at ∼30 nm with a long tail toward smaller distances, as expected for 2D sampling of different molecular orientations in 3D. Fitting the histogram with a model that considers this effect (Method) deduces a distance of 30.8 nm (Fig. 3e, yellow line). Additional analysis and data on another DNA origami can be found in Supplementary Fig. 8.

**Fig. 3.**
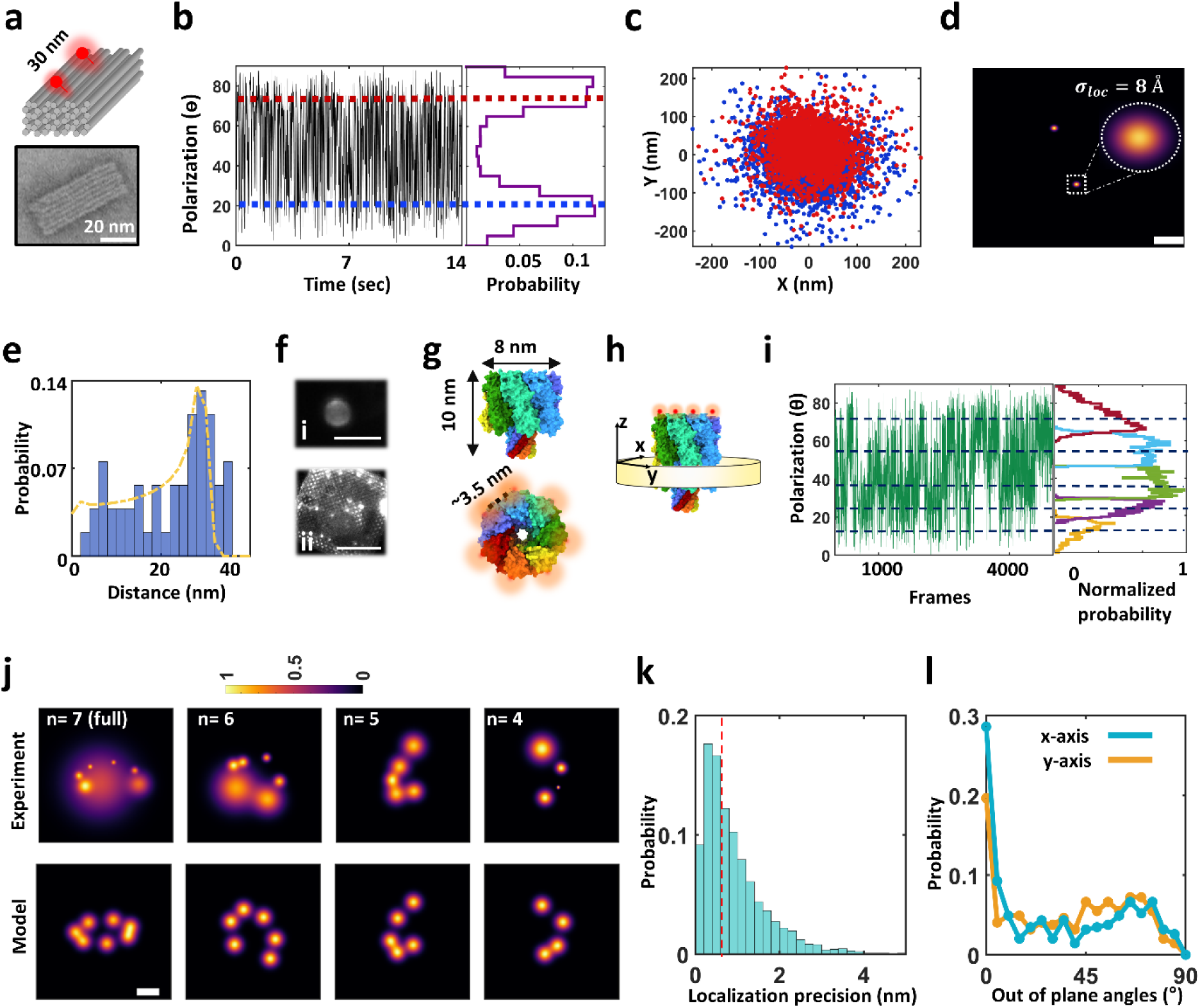
Super-resolution imaging in vitreous ice with Ångstrom precision. **a**, PF3 cuboid DNA origami structure with 2 ATTO647N molecules labelled at separation of 30 nm. Top panel is a schematic, and bottom panel is a TEM image. **b**, Polarization time trace obtained from one PSF of the DNA origami sample. The trace shows two peaks at orientations of ∼ 20° and ∼75° (blue and red lines). We note that this trace is not a continuous time trace, but it rather represents the on-states of the fluorophores during the recording, i.e., off-times were removed (see Fig. 1b as an example). **c**, Scatter plot of the localizations from the polarization events obtained from the two cameras. **d**, By clustering the frames associated with each polarization state (using DISC algorithm, see method section), we generate a 2D resolved image which depicts the position of each fluorophore. The full width at half maximum of the position reflects the localization uncertainty. Scale bar is 10 nm. **e**, Distance histogram obtained from 55 projections with a fit (yellow line) using a model (see Methods). **f**, Vitrified GUVs and SLB imaged in our cryogenic microscope. (i) is a GUV, and (ii) is SLB with ∼0.2 nM fluorescence concentration. Scale bar is 5 μm and 50 μm respectively. **g**, X-ray resolved structure of αHL (PDB: 7ahl), where each protomer is represented with a different color. The protein forms a symmetric heptamer, where the distance between the protomers is ∼ 3.5 nm measured from the last amino acid at the C-terminal side. Red spots in the lower panel indicate the positions of the fluorophores. **h**, axis definition of αHL for protein orientation discussion. **i**, Exemplary polarization time trace from a single PSF shows 5 polarization states. The segmented small histograms indicate the polarization states overt time (see method section). **j** 2D resolved images of single molecules resolved with different examples of 7, 6, 5, and 4 polarization states (top panel). The projection of each were fitted to a model structure generated from PDB:7ahl (bottom panel), revealing a very good agreement within our estimated distance error and localization precision. Scale bar is 3 nm. **k**, Localization precision histogram from the recorded molecules. **l**, Probability of out-of-plane orientation of the particles in the range of [0°-90°], as estimated from a simulated annealing algorithm (Methods).

### Membrane proteins reconstructed in a synthetic lipid membrane

In this section, we benchmark our method by studying membrane proteins reconstructed in a synthetic lipid membrane. We demonstrate the ability of the technique to preserve and image model membranes on the TEM grid. Figure 3f shows that giant unilamellar vesicles (GUVs) and supported lipid bilayer (SLB) could be preserved during the vitrification process and imaged via fluorescence labeling (Methods). To reconstitute proteins into the synthetic lipid membranes, we chose αHL from *Staphylococcus aureus* (Fig. 3g-h) as a model system, which is known to form stable structure of seven protein subunits in the presence of a lipid environment (*57, 58*). We specifically labelled monomeric αHL via an engineered 6x histidine tag located at its C-terminal domain, using Tris-NTA iFlour 647 fluorophore (∼ nM binding affinity, see Methods for further information). The protein was then reconstituted in 200 nm small unilamellar vesicles (SUVs) prepared form 1,2-diphytanoyl-sn-glycero-3-phosphocholine (DPHPC) and incubated with plasma-treated UF TEM grids with 2 nm carbon to form an SLB. The grid was plunge frozen and transferred to our cryogenic microscope (Movie 7). We elaborate on the validation of protein reconstitution in SUVs in the Methods section and Supplementary Figs. 9.

For each detected PSF in the sample, we recorded a polarization time trace and selected particles that had three or more polarization components and maintained a maximum distance below 10 nm. Here, the number of polarization components was determined using unsupervised clustering algorithm (*59*). Figure 3i shows an example with five polarization components. In Fig. 3j, we present several cases of 2D-resolved images from different PSFs, which show fully and partially labeled or assembled protein complexes. Figure 3k plots the distribution of the localization precisions from individual molecules detected on the particles that were included in the analysis. A median localization precision of 7 Å (N=346 molecules) enables us to resolve the full heptameric configuration of αHL, where the smallest distance between protomers is around 3.5 nm (Fig. 3j, n=7). See also Supplementary Fig. 10 for other 2D projections. We fitted the 2D-resolved image to a projection that was model generated based on the known crystal structure (PDB:7ahl)(*57*). The outcome (Fig. 3j, lower panel) indicates a good match between our data and the model within the estimated median distance error of 1.8 nm (Supplementary Fig. 10).

The 2D projection of the fluorophores on a membrane protein allows us to assess the plane in which it is situated. The data in Fig. 3l reveal that, as expected, orientations out of the plane of the membrane are not favored. However, the presence of the tail at severe angles might indicates presence of intact vesicles in the samples, where αHL can freely adopt any 3D random orientation. For example, the full heptamer configuration (Fig. 3j, n=7) is rotated by 3ᵒ about the x-axis and 56ᵒ about the y-axis with respect to a fully flat (top view) structure (Fig. 3h), while the projection with 6 fluorophores (n=6) is rotated by 37ᵒ about the x-axis and 3ᵒ about the y-axis. One can also apply the same algorithm used in Cryo-EM to the 2D projections and calculate the fluorophore configurations in 3D (*43*). However, this procedure is confronted with the challenge that its yield is substantially reduced if the labeling strategy, as is the case for Tris-NTA iFlour 647 fluorophore, is based on affinity rather than a covalent bond. Moreover, resolving a large number of fluorophores in one protein particle requires optimal blinking and polarization behavior from each fluorophore (Supplementary Fig. 10). It turns out that one can get around this difficulty for a structure of known symmetry by selecting under-labeled particles and analyzing the distribution of measured distances. Since successful labeling is more likely under these conditions, one obtains superior statistics.

In the case of symmetric heptamer geometry of side length ∼3.5 nm labeled with only three fluorophores, one expects four different classes with three distinct side lengths of 3.5, 6, 7.5 nm (Fig. 4a). Figure 4b shows a selection of experimentally measured particles with three fluorophores (see supplementary Fig. 11 for additional projections). The histogram in Fig. 4c displays the spread of all measured distances from 179 such particles. A robust assignment of distances is not possible because each side-length component can also appear shorter in a 2D projection, thus broadening its histogram component to lower values and causing overlaps between multiple side-lengths (see supplementary Fig. 12).

**Fig. 4.**
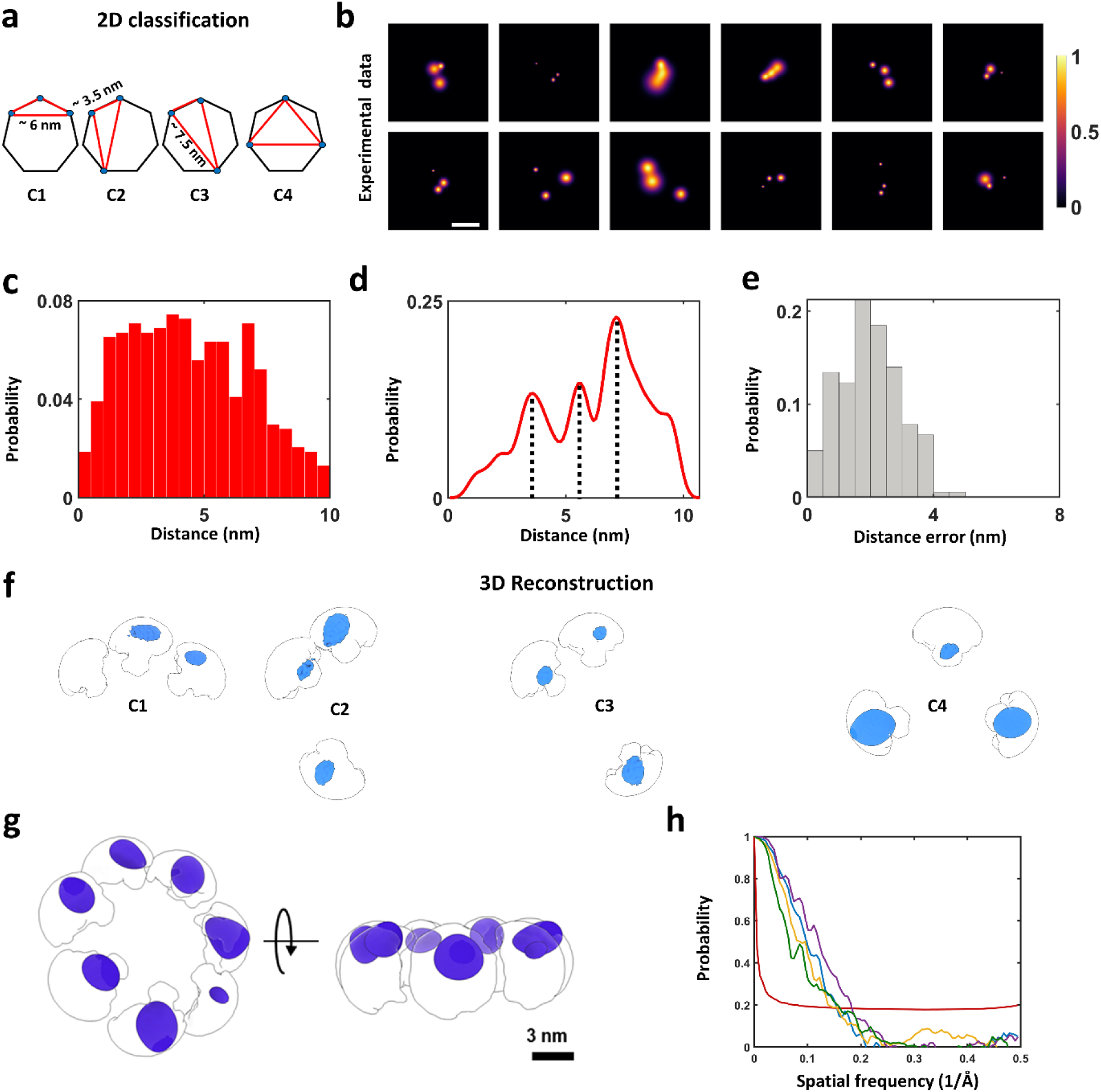
2D classification and 3D reconstruction of αHL. **a**, Heptameric particles labelled with 3 fluorophores sustain triangles that fall into 4 categories, involving side lengths of ∼ 3.5, 6, and 7.5 nm. **b**, Twelve examples of 2D images of single molecules resolved from PSFs with 3 polarization states. Scale bar is 5 nm. **c**, Pair-wise distance histogram calculated from 179 2D projections such as those shown in (b). **d**, Probability histogram of the maximum side-lengths in 2D projections shows clear three peaks at positions that agree very well with the expectations of the model. **e**, Distance errors as calculated by taking all the localization precision of each localization spot (Methods). **f**, Particles were classified into the 4 inherent classes (Supplementary Figs. 11 and 13) in order to determine their 3D reconstructions separately before merging them to form the full structure. The blue spheroids are the outcome of the 3D reconstruction algorithm, which were fitted into the theoretical volume (white clouds) that is accessible to the fluorophores after taking the dye linker into account (*60*). **g**, Final reconstruction of the heptamer model (Movie 8). Purple spheroids are the final 3D reconstructions from all classes. **h**, FSC analysis of the 4 reconstituted classes. Green, yellow, blue and purple curves represent class 1, 2, 3, and 4, respectively. The red curve is the half-bit criteria used to assess the resolution of the 3D reconstruction.

Interestingly, a clever solution can be implemented for the three edges of a triangle that is situated in a randomly-oriented plane, in order to separate multiple side-lengths. In the simplified case of an equilateral triangle of side length *d* regardless of orientation, the 2D projection of one of the triangle’s side lengths will always fall in the range between *d* and 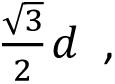 thus, remaining close to *d* (Supplementary Fig. 12). This feature makes the distance *d* become pronounced in the 2D image plane. The situation is more complex for an arbitrary triangle, but the larger side length will always continue to be dominant, accompanied by a smaller tail. Figure 4d presents the outcome of such an analysis when applied to the data included in Fig. 4c, revealing three clear peaks at 3.3, 5.5 and 7 nm, which match the geometry of αHL very well. The deviations in the measured distances are within our median localization precision of 0.7 nm (red line, Fig. 3k), which translates to a median distance uncertainty of 1.8 nm (Fig. 4e). We validated this result also by fitting the data from simulations (Supplementary Fig. 13).

To generate a 3D structural model of αHL from our experimentally resolved 2D projections, a separation of multiple labeling configurations is required to avoid ensemble-averaging. To do so, we subjected the projections of particles with three fluorophores per PSF (Supplementary Fig. 11) to a supervised template classification based on the protein model (Fig. 3g, Supplementary Fig. 13, and Supplementary Method 3). Here, the experimental projections were assigned to each class based on the template matching score, ranging between 0 and 1. A particle was assigned to one of the classes if its score was the highest among the others. Then we selected the particles that scored above 0.8 in each class and reconstructed the 3D localization probability distribution of the fluorophore positions (Fig. 4f). We fitted all the 3D reconstruction models into the theoretically accessible volume of the fluorophores attached to the C-terminal part of the protein and merged the 3D reconstructions together to build the complete heptamer structure of the protein complex (Fig. 4g, Movie 8). By computing the Fourier shell correlation (FSC) of the 3D reconstructions, we obtained a resolution of 6 Å for the reconstituted model (Fig. 4h). Again, fitting the distance distribution of the classified particles separately, indicates a good match with the heptamer model (Supplementary Fig. 13).

### Resolving membrane proteins in their native cellular environment

The procedures described above can also be applied to intact cells and cell membranes. Here too, one can directly benefit from the protocols established in the Cryo-EM field for sample preparation. Given the thin slices that result in a typical realization, background autofluorescence remains manageable, as we reported very recently in two studies on membrane proteins (*61, 62*). In both experiments, we imaged transmembrane proteins in their native plasma membrane following an unroofing procedure (*61, 63*). As depicted in Fig. 5a–c, this process isolates an intact cell membrane on a TEM grid by removing the upper portion of the cells. The sample is then plunge-frozen, transferred, and imaged as described in previous sections. This unroofing procedure was shown to maintain Ca^+2^-dependent exocytosis activity (*63*). More recently, it was implemented in a Cryo-ET imaging pipeline, where it demonstrated retention of high-resolution structural information from proteins close to the cell membrane (*64*).

**Fig. 5.**
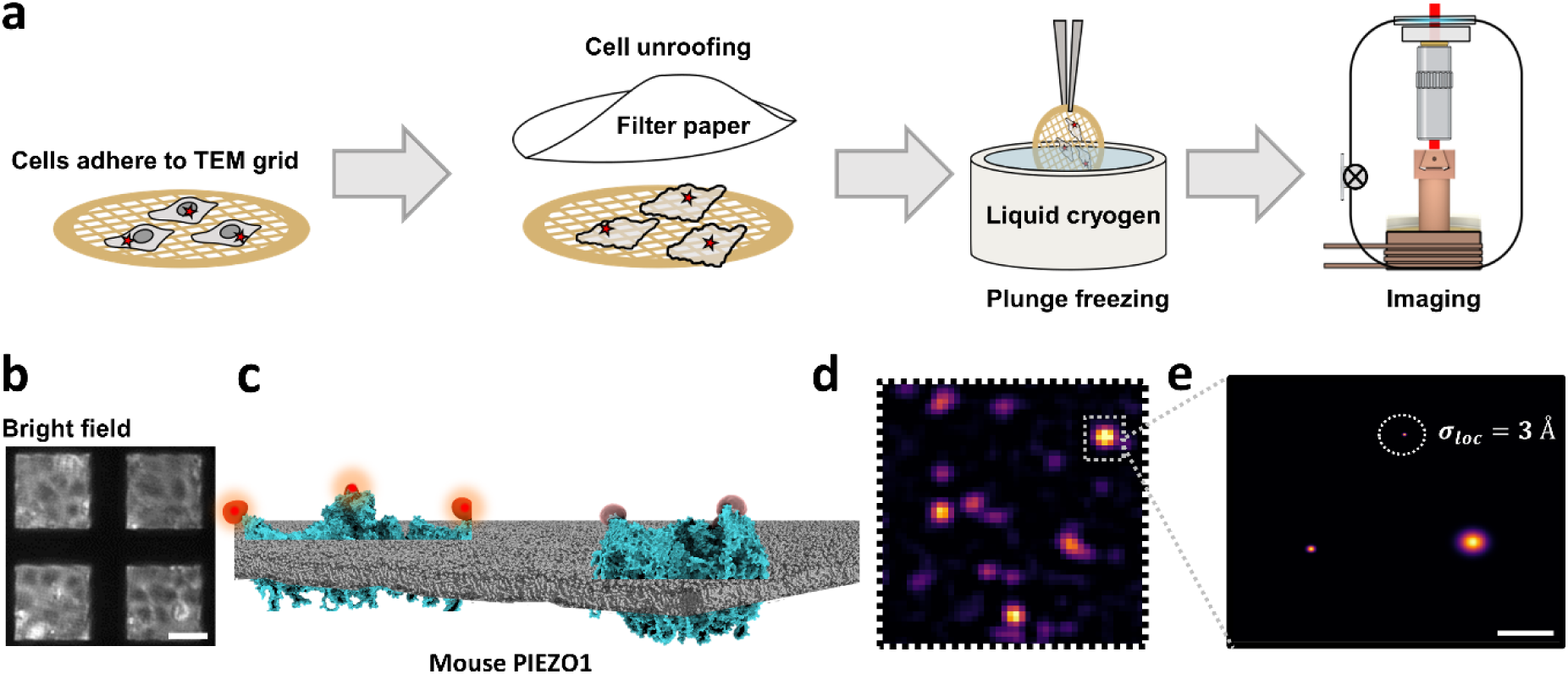
Imaging membrane proteins in their native cellular membrane. **a**, Pipeline of the sample preparation: cells are first unroofing on a TEM grid, plunge frozen and transferred to the microscope cryostat. **b**, Bright-field image of mPIEZO1 transfected COS7 cells on a UF TEM grid before unroofing. Scale bar is 50 µm. **c**, Schematics of fluorescence-labelled mPIEZO1 proteins in a cell membrane. **d**, A fluorescence image (10×10 μm^2^) recorded by our cryogenic optical microscope from one of the mesh squares depicted in (b). The image is a sum of 10000 raw frames**. e**, 2D image generated from a PSF with three polarization states reveals an mPIEZO1 conformation in which the extremities are separated by ∼ 30 nm from each other. Scale bar is 10 nm (*56*).

Figure 5d presents an example of the fluorescence image obtained from the heterologous expressed mouse PIEZO1 protein (mPIEZO1), a mechanosensitive ion channel in COS7 cells. Figure 5e shows an example, where one conformational snapshot of the protein was resolved within one diffraction-limited PSF, in native cellular membrane environment (*62*). In a second study, application of spCryo-LM as discussed in this work allowed us to image the clustering of endogenously expressed muscarinic acetylcholine receptor (M2R), a subfamily of the G protein coupled receptors (GPCRs), in native cardiac derived HL1 cells (*61*).

## Discussion and outlook

By combining super-resolution fluorescence microscopy with shock-freezing, we have established single-particle cryogenic light microscopy (spCryo-LM) as a powerful approach for reaching Ångstrom resolution in localization of specifically-labelled sites of biological samples in their native state. The high photon counts obtained from each fluorescent label under wide-field illumination provides a high yield and, thus, a very robust statistical analysis of single-particle data. For example, a single field of view (80 x 80 μm²) can reach Ångstrom precision from more than 3000 localizations in just 15 minutes. Two concrete recent reports demonstrate the applicability of our method to new discoveries in the conformations and clustering of membrane proteins (*61, 62*). Our general approach of shock-freezing samples and transfer to a cryogenic microscope can also be used with other forms of optical microscopy. In particular, suppression of dephasing in electronic transitions at LHe temperature will offer both larger cross sections and narrower Raman spectra, opening the door to sensitive label-free microscopy.

Other optical methods of current interest such as RESI (*27*) and MINFLUX (*26*) are currently limited to chemically fixated samples. Compared to RESI, our method can observe up to seven copies of the same fluorophore on a protein of interest in a single acquisition, whereas imaging seven subunits in RESI at high resolution would need as many different DNA tags as well as exchange of buffer solution, requiring hours of measurement (*27*). Application of MINFLUX to shock-frozen samples confronts the challenge of inhibitive heating and devitrification by a strong laser beam (*65*).

spCryo-LM also offers some benefits as compared to high-resolution variants of electron microscopy. While the resolution of EM is intrinsically superior to light microscopy, the low electron contrast of biological matter often makes it challenging to resolve different domains of proteins, especially when considering limitations imposed by the acceptable level of electron dose (*10, 14*). The low contrast also brings about the need for averaging over many thousands of single particles and the use of template matching algorithms, e.g., as used in Cryo-ET (*10*). This feature poses a severe limitation for high-resolution investigations of proteins in their complex native environment due to a large background. Combination of electron microscopy with fluorescent labeling is, thus, highly desirable and, indeed, many efforts have already initiated correlative studies (*34, 66*). Being developed for shock-frozen samples on standard TEM grids, our work establishes a reliable pipeline for correlative imaging, promising Ångstrom resolution both in light and electron microscopies at the single particle level.

## Methods

### Cryogenic sample transfer

We describe the whole sample transfer process in the SI, including a step-by-step protocol. All information can be found in Fig 1, Supplementary Fig. 1-4, Supplementary Method 1, and Movie 1.

### Preparation of vitrified fluorescent molecules

In this work, we tested different TEM grids. Carbon TEM grids (R3.5/1, 200 mesh, or other carbon mesh shapes) were plasma cleaned using a Diener Pico 500 W plasma cleaner at 14% plasma power for 10 seconds inside a Faraday cage. Naked UltraAuFoil TEM grids were plasma cleaned at 30% plasma power for 40 seconds, while those with a 2 nm carbon coating were plasma cleaned at 24% plasma power with a 30-second incubation time. For single-molecule fluorescence measurements, 3.5 μl of 50-150 pM ATTO647N-maleimide fluorophores in a 25 mM HEPES buffer solution were loaded onto the TEM grid. The sample was then blotted after a 30-second incubation time using a Vitrobot (IV, Thermofisher) with standard Vitrobot blotting paper (47000-100, PLANO GmbH) and the following parameters: 1-second blotting time,-10 force, 0 wait time, 100% humidity, and 4°C. After plunge-freezing, the sample was either stored at liquid nitrogen temperature or directly transferred into our optical microscope, as described in detail in Supplementary Method 1. Once the sample was transferred into the optical microscope, we allowed it to relax thermally and mechanically from 80 K down to 8 K for approximately 1-2 hours before starting data acquisition.

For the sample preparation procedure for the examination of vitreous ice, please see Supplementary Method 2.

### Vitreous ice determination by cryo-electron microscopy

To examine the quality of vitrified samples (see Supplementary Method 2), we utilized a 300 kV Titan Krios G2 transmission electron microscope (Thermo Fisher Scientific) equipped with a K3 direct electron detector and a Bioquantum energy filter (Gatan). Initially, we created grid atlases of all samples at low magnification (34x) using SerialEM software (67). Next, the grid positions around the center and very far away from the center were imaged at eucentric height at an intermediate magnification (2250x, corresponding to approx. 3.9 nm/px) to assess local ice thickness and hole coverage. High-magnification exposures (105kx, corresponding to 0.85 Å/px) with a total dose of around 60 e/Å² were then recorded at the center of ice-covered holes. We analyzed the power spectra of these high-magnification exposures to detect the presence and intensity of crystalline ice diffraction rings, which are indicative of unwanted ice formation. The characteristic band between 6 and 30 Å in the power spectra can be used to evaluate the presence of crystalline ice (50, 51). Our analysis revealed that only the sample pre-exposed to 1 kW/cm2 exhibited significant devitrification in the exposed region (Fig. 2B and Supplementary Fig. 6). Notably, strong ice formation was also observed at various locations throughout the grid, regardless of the exposure state, likely due to the accumulation of transfer ice during sample preparation steps such as dewar transfer, grid clipping, and microscope transfer. For a detailed discussion on the effect of laser intensity on vitreous ice quality, please refer to Supplementary Method 2 and Supplementary Fig. 6.

### Photophysics characterization

To study the photophysical behavior under different conditions (e.g., temperature and pressure), we opted to image the sample in a polyvinyl alcohol (PVA) polymer matrix. This choice was made for two reasons. First, we checked that the photophysical behavior of ATTO647N embedded in polymer was not significantly different from that in vitreous ice (see Supplementary Fig. 14). Second, imaging in polymer allowed us to measure the very same sample under all conditions, including room temperature, where ice would melt. For this purpose, we prepared a ∼ 10-20 pM solution of each organic dye (ATTO647N, iFluor 647, Cy5, Alexa 647) in 50 mM HEPES buffer containing 1% PVA. The solution was spin coated onto a glass substrate and then inserted into our cryogenic optical microscope (*53*). We first recoded the sample at RT and atmospheric pressure. The chamber was then evacuated to ∼ 10-^4^ mbar, and a different field of view (FOV) of the same sample was recoded. Next, we cooled down the chamber using LHe in combination with a heater on the cold finger, which is controlled via a dedicated temperature controller (Lakeshore) in order to balance the temperature at 77 K. After temperature stabilization (∼ 1 h), we recoded another FOV of the same sample. Finally, we turned off the heater and allowed the system to cool further to LHe temperature ∼ 8K and to relax before acquiring data. In all conditions, the sample was illuminated with a laser intensity of ∼ 0.6 kW/cm^2^ and recorded at a frame rate of 70 Hz. The recorded data were then analyzed using our custom-written code for detecting and clustering the molecules (*53*). The off-on ratio from each PSF/molecule was calculated by dividing the full intensity trajectory (until bleaching point) into 50-frame bin size. For each bin, we calculated the ratio of off-frames to on-frames, then averaged these values across all bins. The final off-on ratio was plotted using a violin plot or as histogram. We note that our imaging frame rate of 70Hz for a FOV of 48 μm x 71 μm limits the temporal resolution of on-times to 14 ms.

### GUV preparation

GUVs were prepared following previously published protocol (*68*). To prepare fluorescently labeled GUV’s, we dissolved 5mg 1,2-Dimyristoyl-sn-glycero-3-phosphoethanolamin – ATTO647N (DMPE-ATTO647N, AD 647N-191, ATTO-TEC) into 500 μl of 20:80 methanol chloroform mixture to obtain stock solutions of 10mg/ml and 0.1 mg/ml. We aliquoted the solution to ∼100 μl volume, and then desiccated them over night at 1e-2 mbar and RT. Then we sealed the tubes and stored them at-20 °C. To prepare the GUV lipid mix, we took 35 µl of 0.1 mg/ml DMPE ATTO-647N and we mixed it with 200 µl matrix lipids 1,2-dioleoyl-sn-glycero-3-phosphocholine (DOPC, 850375 Avanti Polar Lipids) (10mg/ml) and 1765 μl pure chloroform to arrive at a final lipid concentration of 1 mg/ml. DOPC were prepared in a similar way but by mixing the powder with pure chloroform. We then prepared 25 mm coverslips which were pre-washed with 100% ethanol and then plasma cleaned at 100% plasma power for 10 min. We applied 150 μl of 5% poly-vinyl-alcohol (PVA) (8.14894, Sigma Aldrich) and spin coated at 1200 rpm for 30 s. We then dried the sample at 70 °C for ∼ 1.5 h. We applied 5 μl GUV lipid mix and spread the solution across the surface. We then dried the sample via desiccation at 1e-2 mbar for 15-30 min. Next, we added a chamber ring and glued it onto the coverslip with dentist glue. Finally, we added 300 μl aqueous buffer solution and incubated for ∼ 2 h. GUV formation was monitored under standard fluorescence microscope. To image the vitrified GUV’s using our optical microscope, a carbon TEM grid was incubated with 5% PVA and then blotted manually and the solution was allowed to dry. We then applied 3.5 μl of the GUV solution and let it spread on the grid for 30 s, followed by blotting with the following parameters: 3s blotting time,-5 force, 0 wait time at 100% humidity and 4 ^ᵒ^C. Samples were then transferred into our optical microscope as described before. We recorded several images of the grid which showed intact GUV’s, using 645 nm laser at ∼ 0.2 kW/cm^2^ laser intensity and 32 Hz frame rate.

### SUV preparation and SLB formation

Small unilamellar vesicles (SUVs) were prepared following a previously published protocol (*69*). To generate fluorescently labeled SUVs, we mixed 35 or 3.5 μl DMPE-ATTO647N (0.1 mg/ml) with 200 µl DOPC (10 mg/ml) for different fluorophore densities. We dried the solution onto the inner surface of the vial followed by desiccation overnight. We added 2 ml aqueous buffer and vortexed to generate a milky suspension. To obtain homogenous SUVs with well-defined diameter, we extruded the milky suspension sample through a 200 nm pore filter (Whatman Anotop) 37 times. In the case of αHL protein, we produced SUVs from 1,2-diphytanoyl-sn-glycero-3-phosphocholine (850356 Avanti Polar Lipids) following the same procedure. TEM grids with 2nm carbon on top were incubated with SUV solution for 5 min, followed by quick dip in a clean buffer. We applied 3.5 μl clean buffer, blot, and plunge-froze as indicated previously. We used a Teflon sheet facing the side of the membrane and standard blotting paper on the other side. The sample was then transferred and imaged as mentioned previously.

### smURFP purification and imaging

The gene expression smURFP obtained as a gift from Erik Rodriguez & Roger Tsien (Addgene plasmid # 80341; http://n2t.net/addgene:80341; RRID:Addgene_80341) (*55*). One Shot TOP 10 competent cells (C404010 Thermofisher) were transformed with protein vectors and were grown overnight in 10 ml LB + ampicillin. The 10 ml were added to warm 100 ml LB in a 250 ml beaker for large-scale purification. Expression was induced after 2 h incubation by adding 0.2% arabinose, and the cells were incubated at 37 °C overnight at 225 rpm shaking. Following expression, bacteria were harvested and the proteins were purified on a Ni-NTA resin (GE Healthcare) with an elution step involving 250 mM imidazole. This purification was followed by buffer exchange using Amicon® Ultra Centrifugal Filter, 50 kDa to remove imidazole from the solution. The sample purity and protein fluorescence were assessed by native-page and SDS-page gel. The protein was aliquoted and stored at-80 °C. For imaging, UF-2C TME grid were plasma cleaned as mentioned previously. 150 pM of smURFP was applied on a UF-2C TME grid and blotted at-2 force. The sample was then transferred and imaged with laser intensity at 0.3 KW/cm^2^ and 70Hz frame rate.

### DNA nanoruler imaging

30 nm DNA nanoruler was purchased from Tilbit nanosytems GmbH, and 116 nm was purchased from GATTAquant GmbH. In both cases, ATTO647N was used as label. For imaging, UF-2C TME grid were plasma cleaned at 15% power for 25 s in a Faraday cage. DNA sample was diluted by a factor of 4 from the stock sample as received from the manufacture, in a buffer contacting 20 mM MgCl_2_. 3.5 μl of the sample was applied onto the TEM grid and blotted at-5 force for 2 s. The sample was transferred and imaged with laser intensity at 0.6 KW/cm^2^ and 50-70Hz frame rate.

### Alpha-hemolysin labeling and lipid reconstitution

C-terminal His-tagged αHL protein from *Staphylococcus aureus* was purchased from Hölzel Diagnostika Handels GmbH (catalogue number CSB-YP639324FLFc7-200). The protein was dissolved in 25 mM HEPES buffer, 25 mM KCl at pH 8 to a concentration of 10 µM. The sample was next desalted using size-exclusion column PD SpinTrap G-25 (Cytiva, 28918004) to remove older buffer residuals. The protein was specifically labeled via the histidine linker on the C-terminal side of the protein with the dye HIS Lite™ iFluor™ 647 Tris NTA-Ni Complex which was purchased from AAT Bioquest (catalogue number 12618). The protein was reacted with dyes at a ratio of 1:1 respectively for 2 h at RT. The protein was then desalted from the excess of dyes using the same desalting column. The labeling efficiency was estimated using an absorption spectrometer (Nanodrop 2000, ThermoFischer), confirming ∼ 100% labeling efficiency. However, as the sample is diluted and washed further during the sample preparation, the labeling efficiency is expected to decrease since the labeling is not covalent. The sample was aliquoted, flash frozen and stored at-80 °C. To generate the protein lipid construct, we followed previously published protocols (*70*). First, we incubated the protein with a pre-made 200 nm SUVs at 38 °C for 1-2 hr. Then we performed size exclusion to purify the SUVs only and remove unbound free protein. We validate the incorporation of αHL into vesicles via correlative interferometric scattering (iSCAT) and florescence imaging (see Supplementary Fig. 9). We incubated plasma-cleaned UF TEM grid with 2 nm carbon on top with the protein-lipid mixture for 5 min at RT. The sample was then dipped several times in clean buffer before plunge-freezing. In the plunge-freezing process the sample was blotted with a teflon paper on the side facing the lipids and standard paper on the other side. We note that some residual SUVs might still be present on the sample as the washing was very gentle. In addition, as the concentration of αHL is lower than that at the labeling stage, some of the fluorophores might dissociate as the binding of the fluorophore to the 6x Histidine tag is in the nM range.

### Optical setup

Our experiments were conducted in a custom-built cryogenic microscope, which utilizes a Janis ST-500 flow cryostat to maintain liquid helium temperatures. To facilitate the insertion and removal of vitrified samples, we modified the microscope’s cold finger to accommodate a dovetail-shaped sample cartridge, allowing for high-vacuum and low-temperature operation (see details in Supplementary Figs. 1-4). The mechanical drift of the microscope was measured to be approximately 5 nm/min on average. Additionally, the temperature at the cold finger was verified to be stable at 8.23 K, regardless of the laser intensity used. The temperature characterization of our sample cartridge is presented in Supplementary Figs. 2-3 and described in detail in the supporting information. The optical setup is identical to that used in our previous work, as described in Refs. (*43, 53*). In brief, samples are loaded onto the cold finger and imaged using a 0.90 NA objective lens (MPLAN 100x, Mitutoyo) mounted in vacuum, which focuses the fluorescence onto two separate EMCCD cameras (Andor iXon) in a polarization-resolved configuration. To achieve fast acquisition rates, the field of view was set to 211 x 313 pixels, with a pixel size of 227 nm. The laser intensity was set to approximately 0.65 kW/cm^2^, and images were recorded with an exposure time of 14 ms, which is more than five times faster than the typical off-time. This allows us to capture individual fluorescence bursts and minimize the likelihood of overlapping emission from multiple fluorophores in a single frame. For each field of view, we collected a total of 50,000 to 100,000 frames, depending on the specific experiment

### Image analysis

We employed custom-written MATLAB software, as described in our previous works (*43, 53*), to analyze the raw image stacks from two polarization channels. To further investigate the polarization time-traces of multi-fluorophore particles, we utilized a routine based on the DISC algorithm (*59*). This approach enables us to determine the number of polarization states per point spread function (PSF), which corresponds to the number of fluorophores per PSF or protein particle. The underlying principle is that the dipole orientation of each fluorophore at 8K is random but fixed (*43*), allowing us to annotate each fluorophore over time and localize it with high precision beyond the diffraction limit through coordinate clustering (*43*). By assigning a 2D Gaussian function to each localized polarization/fluorophore, with a width determined by the respective localization precision, we reconstructed 2D super-resolved images. These images provide different projections of the protein molecules within the sample. The 2D images were then subjected to further analysis, including distance measurements, classification, and 3D reconstruction, as outlined below.

### Single-molecule 2D image analysis of αHL

To classify the experimental 2D maps of the αHL, we followed a similar approach as reported in Ref. (*43*). Here particles with three fluorophores inherently generate 4 different classes (see Figure 4a). We simulated multiple 2D projections of the protein spanning all possible 3D orientations using the theoretical structural model (PDB: 7ahl) for each class separately. We then performed 2D cross-correlation of the experimental and simulated maps of all classes. The correlation score, which ranges from 0 to 1 with the latter indicating a 100% match, was used to assign a class to each experimental image. In this case, an image is assigned to a certain class if the correlation score of this class is the highest score among the others. See Supplementary Fig. 12-13 for additional analysis.

### Distance error estimation

The error on the measured distances were calculated in an error propagation manner following 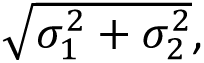 where σ_1,2_ indicate the localization precision of positions 1 and 2, respectively. The latter quantities are calculated as 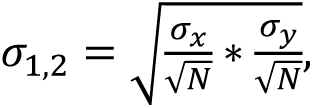 where σ*_x,y_* is the standard deviation of the position in the x and y directions, respectively, and *N* is the number of localizations per position.

### Distance fitting

For more quantitative analysis of the pairwise distances, we used a model described in (*53, 71*). In short, we convolve the Rician distribution of the distance between two spots with finite localization uncertainty, considering the projection onto the image plane to be a cosine function.

### Estimation of particle orientation

To determine the orientation of our particles in 2D space, we utilized a simulated annealing algorithm. This approach employed a triangular model with side lengths of 9, 19, and 34 nm, which was allowed to explore all possible rotation angles, including yaw (z-axis), pitch (x-axis), and roll (y-axis). The goal was to find the 2D projection that maximized the similarity between the projected and experimental images, as measured by 2D cross-correlation (2DCC). To prioritize smaller pitch and roll angles (out-of-plane orientations) while permitting free rotation in yaw (in-plane orientation), we introduced an angular penalty term into the objective function, which took the form:

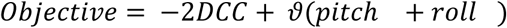

where ϑ represents the penalty weight. Following the estimation of particle orientation, we calculated the probability distribution of angles along the x, y, and z axes

### Tomography and 3D reconstruction

We selected particles that exhibited a high degree of similarity to the simulated 2D projections, as evidenced by a cross-correlation score greater than 0.9 and a localization precision better than 2 nm. The 2D maps of the particles were then normalized and the median of the full width at half-maxima was taken as the localization uncertainty for all particles, in order to obtain a spherical volume. The resulting 2D maps of the αHL protein were constructed on a 120×120 grid with a pixel size of 1.5 Å. The 2D projections were then used as input for the 3D reconstruction subspaceEM algorithm (*72*), with an elliptical Gaussian serving as an unbiased initial structural model. We ran the algorithm for 100 iterations using default settings. The 3D volumes were then refined by fitting them to crystal structures and maps of the dye location (*60*), and edited using ChimeraX (*73*). To assess the resolution of the 3D reconstructions, we computed the Fourier shell correlation (FSC) curves using the FSC server provided by the Protein Data Bank in Europe (https://www.ebi.ac.uk/pdbe/emdb/validation/fsc/). This involved dividing the 2D image data into two equal sets, reconstructing the 3D volumes independently using the previously determined 3D volume as an initial model, and then aligning the two reconstituted volumes to calculate the FSC. The resolution was determined based on the half-bit criterion (*74*), which provides a reliable estimate of the achievable resolution.

### Data visualization and analysis

All data analysis done using MATALB Mathworks software. Some of the figures were plotted using OriginPro 2020. Protein structures and 3D reconstitution volumes were processed using PyMOL v2.4 and ChimeraX v1.7.1.

## Data availability

All data supporting the article are provided in main text and supplementary information. Protein structures were obtained from RCSB Protein Data Bank (7ahl). 3D reconstructions are provided as a source data. Source data are provided with the paper.

## Code availability

The single molecule analysis code was developed by a previous member of our group and published in Ref (*53*). The code for polarization trace fitting is available at DISC. The code for the 3D reconstruction is available from the subspaceEM algorithm (*72*). FSC was calculated using the online tool available at EMBL website https://www.ebi.ac.uk/pdbe/emdb/validation/fsc/.

## Acknowledgements

We thank Daniel Böning and Sebastian Tacke for helpful discussions in the planning phase of the transfer shuttle. Robert Branscheid and Erdmann Spiecker for the generous loan of the Vitrobot shock-freezing tool in our lab, as well as assistance in imaging samples using an electron microscope. We thank Simone Ihloff for help in the cell culture lab, Tobias Utikal for assistance with cryogenics, Martin Blessing for help in generating GUV samples and SUVs, Maksim Schwab for his insights and help with custom engineering of various mechanical part as well as Oliver Bittel for support in establishing the temperature sensors. We also thank Shuan Jiang, Morgan Miller and Hannarae Lee for the help with recording correlative iSCAT-fluorescence data of the αHL sample. In addition, we thank David Albrecht and Morgan Miller for valuable comments on our manuscript.

## Author contributions

F.F.W, H.M. and V.S. conceived the apparatus. H.M. and F.F.W. assembled and characterized the high-vacuum shuttle transfer system and the optical microscope. H.M. prepared, performed and analyzed all the experiments reported in this paper, except for the EM part which were done by D.B. The manuscript was written by V.S. and H.M. The measurements were conceived by H.M. and V.S. The project was supervised by V.S.

## Supplementary Materials

### Supplementary Methods

#### 1. Cryogenic high-vacuum transfer

The design features of the transfer system are primarily dictated by temperature and contamination requirements. Specifically, the plunge-frozen grid must be maintained below 130 K and protected from direct exposure to air. In traditional Cryo-EM holders, the grid is loaded under liquid nitrogen and enclosed by a cold shield as it briefly traverses through air before insertion into the microscope. Residual contamination in the form of small ice crystals is sometimes observed in this process. A direct transition from liquid nitrogen to vacuum has proven to significantly reduce contamination, a critical consideration in correlative studies involving multiple transfers of a grid between different instruments (*1*). We adapted this strategy in our cryogenic optical microscope (*2, 3*) with a series of modifications as depicted in supplementary Figs. 1-4.

First, plunge-frozen grids are mounted on a purpose-built cartridge (supplementary Fig. 1) inside a dedicated preparation chamber as depicted in supplementary Fig. 2a. The preparation chamber includes a working platform that holds a cold stage (cartridge holder) and a TEM grid box groove. A glass vessel (SCH 9, KGW Isotherm, ∼ 120 ml) serves as a liquid nitrogen (LN) container, and a vertical vacuum manipulator is used to lower or raise the glass vessel. The chamber is initially purged with dry nitrogen to reduce water vapor. The glass vessel is then filled with LN and is moved upward to cover the working platform completely. In this case the TEM grid can be picked up and mounted safely onto the sample cartridge in LN. The grid is securely held in place by a clamp with a thin profile to allow positioning it under a long working distance microscope objective (Mitutoyo Plan Apo HR, 100x, 0.9 NA) in the cryostat. To ensure proper surface contact between the sample cartridge and the cold stage, we used two repelling magnets press-fitted onto the both components (see supplementary Fig. 1 a-b)

The high-vacuum transfer shuttle (Pfeiffer, 420MDM040-0500) consisted of a linear manipulator and a cooling stage as core components (supplementary Fig. 2e). The linear manipulator with 500 mm range and maximum torque of 2.3 Nm is connected to a conical shaped copper head that allows sample cartridge loading via a small screw. The high-vacuum shuttle is evacuated to ∼10^-6^ mbar and cooled to LN temperature. The cold stage is actively cooled during the whole transfer process, maintaining a stable temperature below the devitrification point (supplementary Fig. 2g). High-vacuum is maintained within the shuttle by the cold stage after closing the gate valve to the pump. The shuttle interfaces seamlessly with the microscope through a KF50 flange. The external interfaces are flexible such that the design can easily be adapted to suit other microscopes.

Upon loading the TEM-grid we lower the LN vessel to provide side access to the cartridge and begin evacuating the chamber. As some LN remains in the glass vessel, the vacuum level reaches 10^0^-10^-1^ mbar. However, we minimize condensation by extensively purging the preparation chamber with dry nitrogen throughout the entire process until evacuation begins. This significantly reduces the water vapor level in the chamber. Next, we open the high-vacuum shuttle’s gate valve and use the manipulator to load the sample cartridge. Once loaded, the manipulator is retracted and parked on the cold stage to maintain the sample cartridge at LN temperature. Temperature readings of the cartridge before and after the loading step remain stable close to LN temperature (supplementary Fig. 2b,d).

Then, we close the preparation chamber gate valve and allow the transfer shuttle to establish high vacuum level again (10^-6^ mbar). After reestablishing high vacuum in the transfer shuttle, we close its gate valve and vent the connection interface to disconnect it from the preparation chamber. The transfer shuttle is then connected to the cryogenic optical microscope. After establishing high vacuum level (10^-6^ mbar) at the connection interface, we insert the sample cartridge into the microscope’s cold finger. The optical microscope is pre-evacuated to 10^-6^ mbar and precooled to LHe temperature prior to sample insertion. After inserting the sample into the microscope, thermal contact with the cold stage adapter is sustained through the side faces of the dovetailed piece, facilitated by two pairs of repelling magnets embedded in the cartridge and the cold stage adapter. Pogo pins connect the temperature sensor of the sample cartridge to the controller (supplementary Fig. 3). Temperature readings from silicon diodes on the cartridge and the cryostat show 8.3 K and 4.3 K, respectively (supplementary Fig. 3).

The optical microscope (supplementary Fig. 3) is built around a Janis-500 cryostat, which is connected to an extended cold finger with a complementary shape to the sample cartridge. The objective is mounted from the top and has a working distance of 2 mm and numerical aperture (NA) of 0.9. The objective can be positioned against the sample with an x, y, z piezoelectric scan stage. The laser beam is guided through a window to reach the objective and is aligned at the center of the TEM grid (supplementary Fig. 3). We have tested the stability of the optical microscope and found a mechanical drift in the order of ∼ 5 nm/min (supplementary Fig. 3f). See supplementary Fig. 3e for the complete optical scheme of the microscope.

See supplementary protocol for a step-by-step procedure.

#### 2. Validation of vitreous ice stability upon laser illumination

To obtain a sufficiently large number of photons during the on-time blinking of a single fluorophore, it is desirable to excite the molecule at a high rate, leading to laser intensities in the range of 0.3 - 1 KW/cm^2^. However, such high light intensities are known to cause severe sample devitrification at 77 K, especially in the case of carbon mesh TEM grids (*4, 5*). Gold TEM grids were found to handle high optical intensities (∼ 0.6 kW/cm^2^), to possess superior mechanical stability and to maintain stable vitreous ice condition (*4, 6*). To verify this experimentally, we vitrified aqueous samples on several UltrAufoil R2/2 200 mesh (S373-7-UAUF) TEM grids and exposed each one of them to varying laser intensities in the cryostat for few hours. The samples were subsequently retracted from the cryostat and investigated by Cryo-EM imaging (Supplementary Fig. 6A) to examine the presence of devitrified ice.

For this, we used a vitrified fluorescent aqueous solution because in the case of pure water, the holes would appear dark, making it impossible to distinguish the presence of water using an optical microscope. Thus, it would not be easy to asses if the grids were over-blotted. In the case of a fluorescent solution, the signal should be visible across the grid, especially in the empty holes (see Supplementary Fig. 6B).

We used TEM grids made of gold (UltrAufoil R2/2 200 mesh, S373-7-UAUF), which were plasma-cleaned at 24% plasma power for 45 seconds in a Faraday cage (Diener, Pico 500 W). A 3.5 µL of 0.5-1 nM ATTO647N solution was applied to the grid and allowed to spread for 30 seconds. The sample was then blotted using the following parameters: 3 seconds blotting time,-5 blotting force, and 0 waiting time. Next, the sample was transferred into our optical microscope following the procedure outlined in the previous section and was allowed to equilibrate for 1–2 hours.

The samples were illuminated with a 645 nm laser source in wide-field mode at different laser intensities (0, 0.16, 0.34, 0.65, 1 kW/cm²). First, we quickly screened the grid area (each field was exposed for ∼5 seconds) until we reached the center of the TEM grid. We then illuminated each area around the center intensively for 8–15 minutes, corresponding to a typical single-molecule acquisition time discussed in the main manuscript. We used the internal marker on the TEM grid to navigate across the grid. Our field of view was 512 x 512 pixels, with a pixel size of 227 nm, which corresponds to a complete grid mesh size of ∼100 µm x 100 µm (see Supplementary Fig. 6B). In this case, we illuminated a 5 x 5 region around the center of the grid, while the rest was illuminated for a short exposure time of 0.5–1 minute. The fluorescence signal shown in Supplementary Fig. 6B was used to optically confirm the presence of vitreous ice on the sample. The bright spots/regions indicate the presence of the fluorescent solution on the grid, serving as a validation before the subsequent Cryo-EM measurement. Each laser intensity was measured in duplicates/triplicates (2–3 grids each). The whole experiment spanned ∼4 days.

At the end of the measurements, the samples were retracted and immediately stored in a LN storage dewar. They were then transferred to the Cryo-EM facility in Munich for vitreous ice assessment and analyzed as explained in Methods.

All grids illuminated below ∼ 0.7 kW/cm^2^ laser intensity survived and showed vitreous ice conditions, except for one sample (sample number 7), which exhibited crystalline ice patterns (see Supplementary Fig. 6c). We attribute this to bad handling of this particular sample during a transfer process.

#### 3. Particle classification

We used a supervised classification approach for particle classification. To do so, we generated a library of images based on the structural model of αHL (PDB: 7hal). For each of the four classes we rotated the particles in 3D and then projected the image onto the 2D plane. We used a rotation increment of 5° for x, y and z axes. Next, we cross-correlated the experimental image to all the library generated from all the classes, and determined the maximum cross-correlation value (score), ranging from 0 to 1. The class that yielded the highest score is assigned to the experimental image.

To test the accuracy of this approach, we generated random images from all four classes. Here, we used the center position of the fluorophore exactly as in the structural model but with a different localization precision. Then we subjected these images to 2D cross-correlation as explained above. As depicted in supplementary Fig. 13a, if we assume high localization precision 7 Å (similar to our experimental data), the classification reaches 100% accuracy. This also holds for 1 nm localization precision (supplementary Fig. 13b). However, if we increase the localization precision to ∼ 2 nm, ∼ 20% of class 1 and class 2 are misclassified (supplementary Fig. 13c). Next, we simulated data with random centre position displacement of ± 1 nm, assuming 7 Å localization precision. Analysis of particle classification under these conditions revealed substantially increased misclassification rates, particularly for classes with shorter distances such as class 1 and class 2 (supplementary Fig. 13d). Thus, our conclusion is that the lower the localization precision, the larger the misclassification. As a result, if one seeks to obtain informative data for structural biology, one must reach localization precision better than 2 nm.

## Supplementary figures

**Supplementary Fig. 1.**
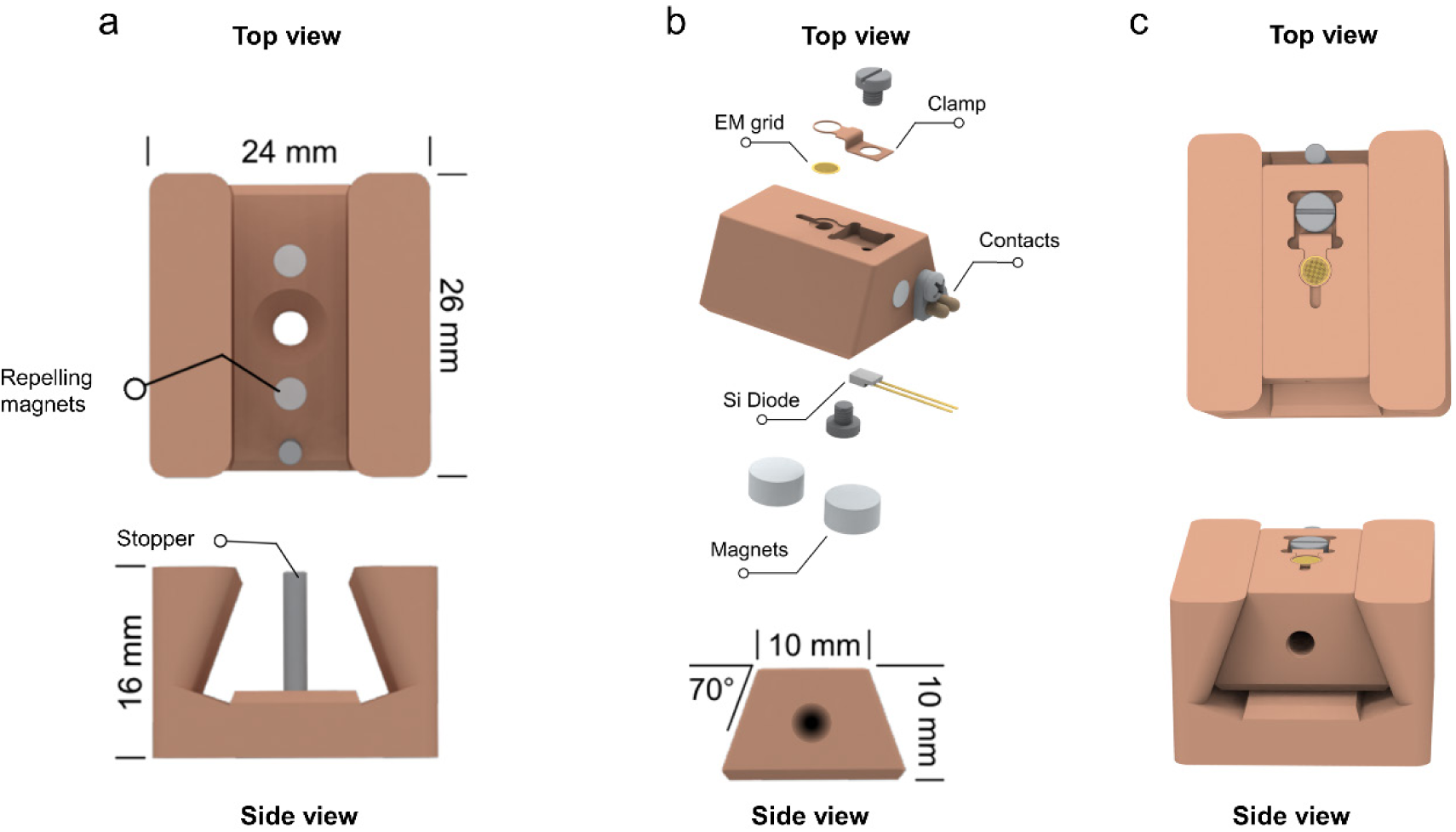
Sample cartridge and cold stage adapter. **a**, Top and side views of the cold stage, designed with a complementary dovetail-shaped structure. It contains two repelling magnets and a stopper to properly position the sample cartridge. **b**, View of the sample cartridge. A single TEM grid is mounted with a small clamp and a screw. The dovetail-shaped cartridge inserts into the cold stage adapter and is held in place by two pairs of repelling magnets and an end-stop. A bare silicon diode temperature sensor (DT670) is integrated into the bottom of the cartridge. **c**, Final assembly of the parts in (a) and (b). The cold stage in (b) is similar to that used in the optical microscope but includes two additional pogo-pins for temperature readout from the sensor integrated into the sample cartridge, see supplementary Fig. 3.

**Supplementary Fig. 2.**
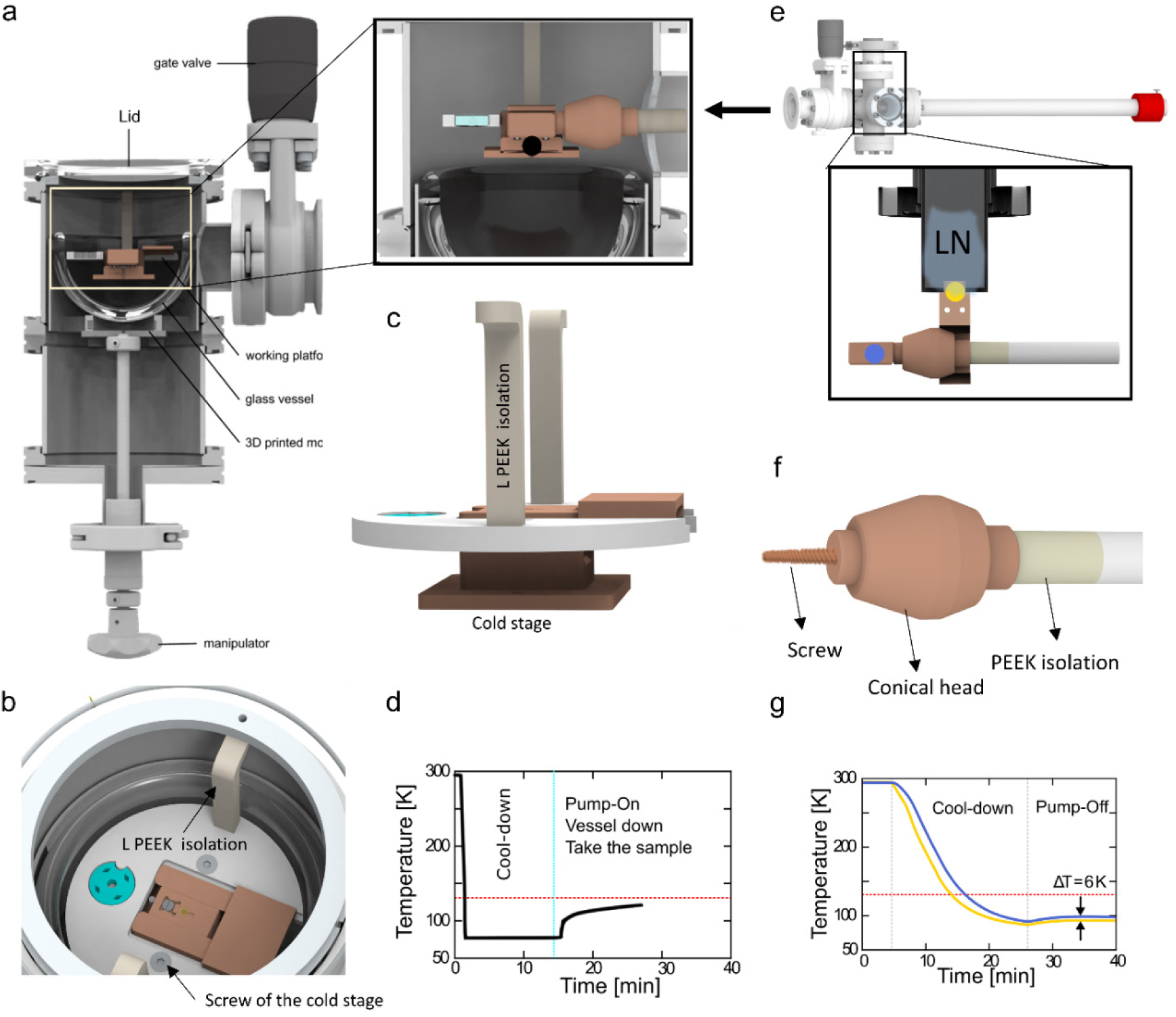
**Sample loading chamber**. **a**, Cut view of the sample loading chamber. An insulated glass vessel (KGW Isotherm) holding up to 120 ml of liquid nitrogen is attached to a vacuum manipulator for vertical movement. In the fully lifted position, the vessel allows complete immersion of the working platform in liquid nitrogen. The TEM grid can be mounted onto the cartridge with a pre-cooled tweezer without the risk of de-vitrification or contamination. The loading chamber is sealed on top with an acrylic plate, connected to a nitrogen gas supply line in the back, and to a pumping station (Pfeiffer) via a gate valve on the side. **Inset**, When the vessel is fully retracted, the transfer manipulator can access the cartridge from the side. **b**, Shows the working platform hanging from two polyether ether ketone (PEEK) isolation L-pieces mounted on the top flange. **c**, Shows a side view of the working platform, featuring a copper cold stage suspended from the platform by two mounting screws **d**, Temperature readout of the sensor placed on the cold stage (black circle, inset of (a)) during the transfer process as describe in supplementary Note 1. The temperature remains below the devitrification point. **e**, High-vacuum shuttle after loading and transfer of the sample cartridge, where the cartridge brought in contact with the cold stage to keep the sample below the devitrification point. **f**, Linear manipulator with the large-mass conical head. The tip of the manipulator is extended with 20 mm poly ether ketone (PEEK) isolation. The conical head contain a screw for loading the sample cartridge via its thread (see supplementary Fig. 1b). **g**, The temperature readouts of the sample cartridge (blue circle in c) and the cold stage (yellow circle in c). The temperature is stable and remains below the devitrification point. Color code is similar to the sensor in c.

**Supplementary Fig. 3.**
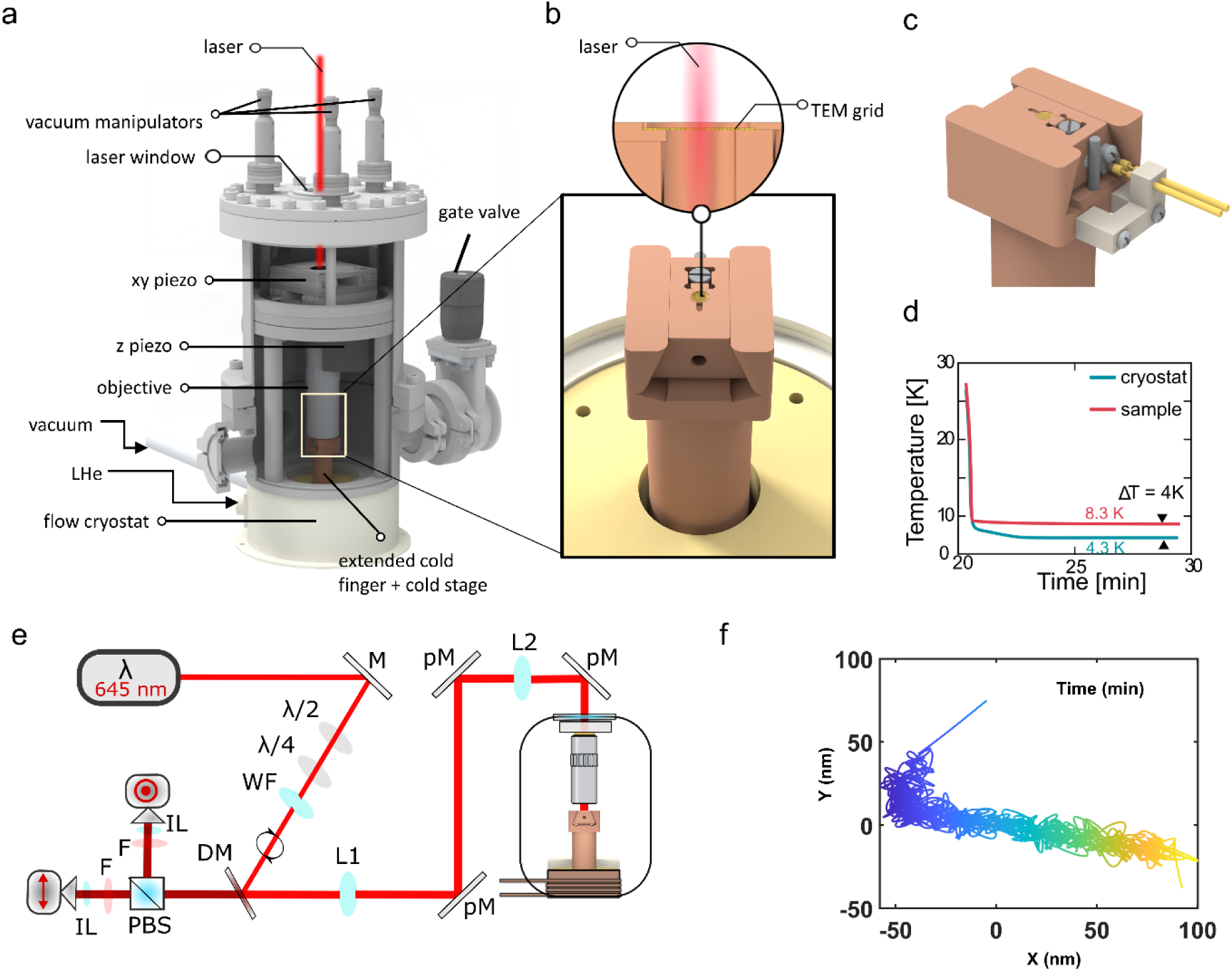
Modified high-NA cryogenic microscope. **a**, View of the cryostat and vacuum system of the optical microscope. The microscope is built around a liquid helium cryostat (Janis-500) shown in the lower part. An extension chamber with four KF40 ports allows access from the side via a gate valve. The cryostat cold finger is extended to place the cold stage adapter on the axis of the vacuum port for inserting the cartridge. An objective (Mitutoyo Plan Apo HR, 100x, 0.9 NA) connected to an x-y-z piezo stage allows sample scanning and focusing. **b**, Close-up and side views of the sample cartridge loaded into the cold stage adapter inside the microscope. The view is from the side of the transfer shuttle access and the gate valve indicated in (a). Inset shows a side view of the optical axis. The laser beam traverses the sample through a hole in the cartridge. **c**, Rear view of the cold stage of the image in (b), depicting the connection between the sample cartridge and the pogo pins assembled on the cold stage for temperature readout. **d**, Temperature readout indicates the sample cartridge stabilized at ∼ 8 K. **e**, Schematics of the optical microscope pathway, which allows us to select the polarization of the fluorophores in our samples as explained in detail in Ref (*2*). A linearly polarized laser beam (645 nm) is transformed to a circularly polarized beam using λ/2 and λ/4 waveplates. A wide-field (WF) lens is inserted to focus the laser beam onto the back focal plane of the objective. L1 and L2 establish a telecentric lens system with 400 mm focal length. The polarization of the laser is maintained along the path using polarization-maintaining coated mirrors (pM). The fluoresce emission is split using a dichroic mirror (DM), and the polarization is split using a polarized beam splitter (PBS). The emission is then filtered using an emission filter (F) and focused onto an EMCCD camera using an imaging lens (IL). The microscope is evacuated up 1×10^-6^ mbar and the cold stage is cooled down to 4 K using liquid Helium. Once the sample inserted, we turn off the vacuum pump and allow the system to relax for 1-2 h before we start acquiring data. **f**, The mechanical stability of our microscope is very high, as we achieve a drift of ∼ 5 nm/min, close to the manufacturer value of ∼ 2 nm/min.

**Supplementary Fig. 4.**
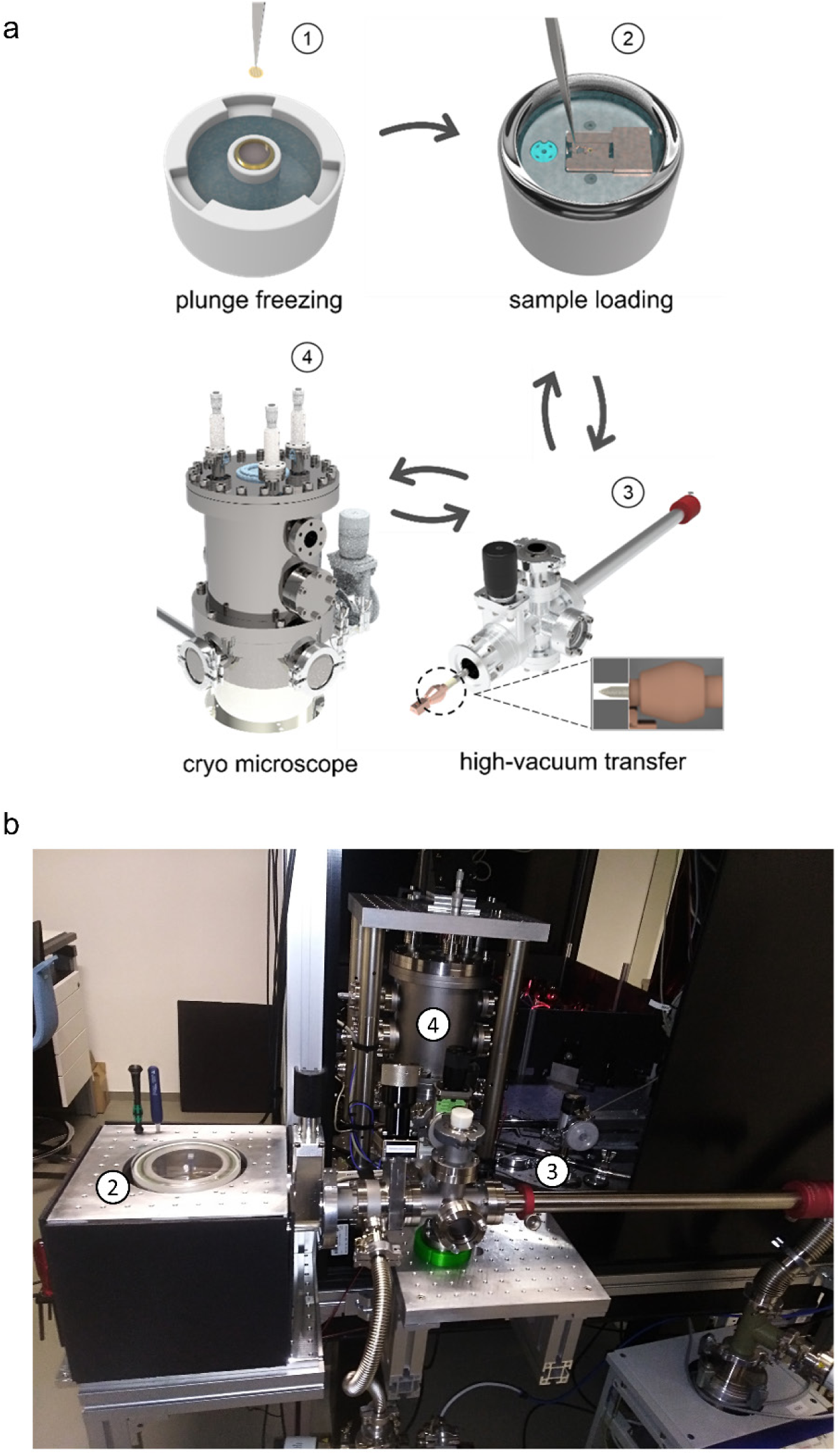
Cryogenic high-vacuum transport shuttle. **a**, Overview of the sample transfer scheme. The sample is plunged into liquid ethane (1) and then loaded onto the sample cartridge inside a liquid nitrogen LN bath in a special preparation chamber (2). A high-vacuum transfer shuttle (3) is used to move the sample from the preparation chamber to the optical microscope (4). This process can also be executed in a reverse order for correlative studies. **b**, Real image of the setup showing the preparation chamber (2), high-vacuum transfer shuttle (3), and the optical microscope (4).

**Supplementary Fig. 5.**
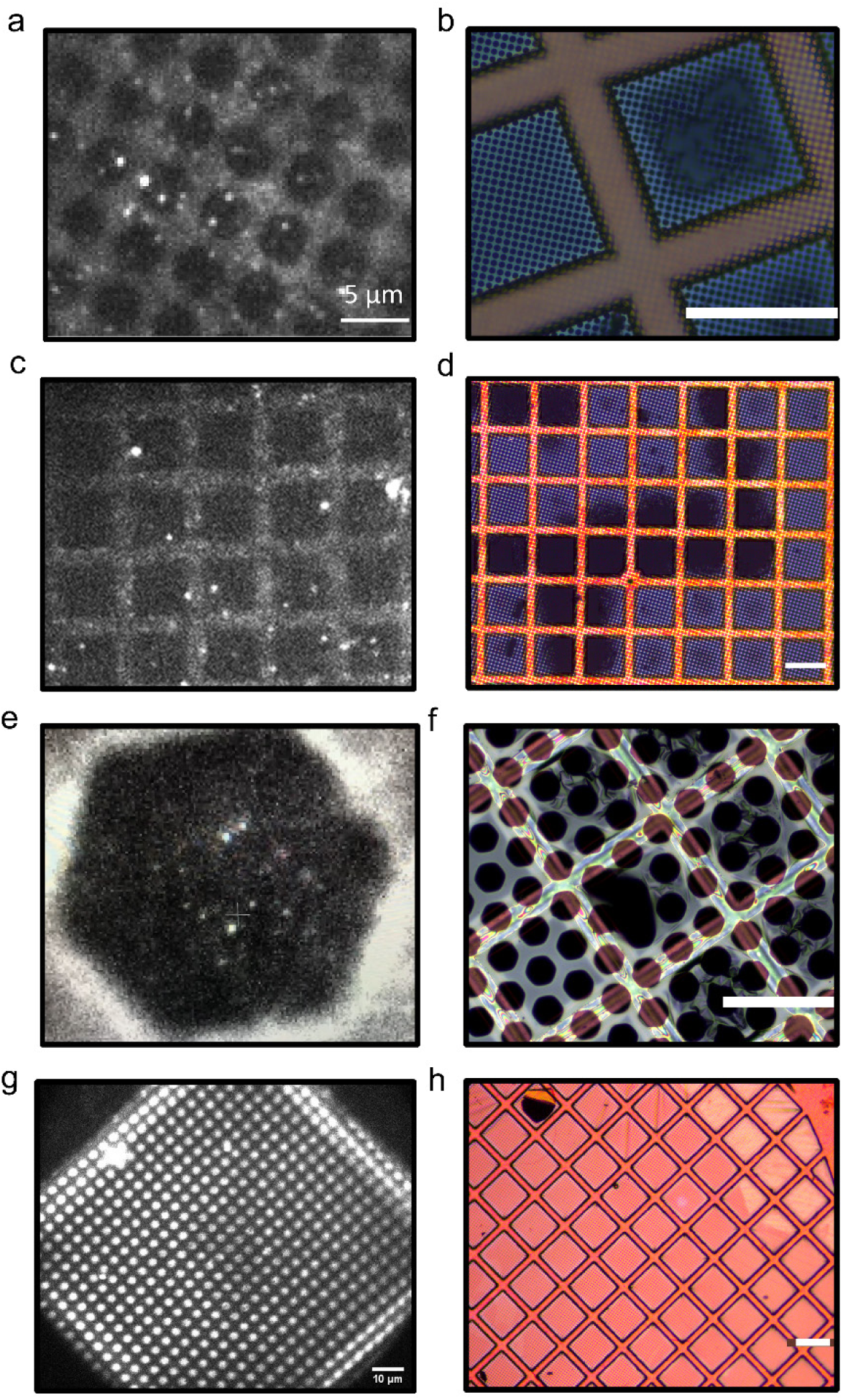
TEM grid mesh stability. Aqueous solutions (25 mM HEPES and 150 pM ATTO647N) were vitrified on different carbon mesh grids TEM grids. The samples were loaded into the microscope and illuminated with a laser intensity of ∼ 0.6 kW/cm^2^ for fluorescence images recordings. Afterwards, the sample was retracted and inspected visually using a bright-field microscope. **a**, Fluorescence image of a carbon mesh grid (Gold, R3.5 200 mesh, S177-7 Plano-GmbH). Bright spots are single fluorescent molecules, and the low intensity grey background is the carbon mesh. **b**, Bright-filed inspection after the recording. The data show that some mesh got damaged. Brown bars are the gold framework, and the bluish mesh is the carbon mesh. **c-d**, Similar to (a-b), but for square carbon mesh grid (Gold, R7/2 square 200 mesh, S117-7 Plano-GmbH). **e-f**, Similar to (a-b) but for hexagonal carbon mesh grid (Gold, R25/15 hexagonal 200 mesh, Hex15-7 Plano-GmbH). The image in (e) shows one hexagonal hole (black region) filled with single fluorescent molecules (bright spots). The carbon support is shown on the periphery with high intensity due to high reflection. **g-h**, Similar to (a-b) but for gold mesh TEM grid (UltrAuFoil 200 mesh, QF R2/2 + 2nm carbon film on top, S373-7-UAUF-2C, Plano GmbH), which is the only grid that remained intact and stable. The bright regions in (g) are holes, containing vitrified aqueous solution with nM fluorescent molecules. The black/grey mesh is the gold mesh support. The scale bar in all images is 100 µm. (see associated Movie 1-3).

**Supplementary Fig. 6.**
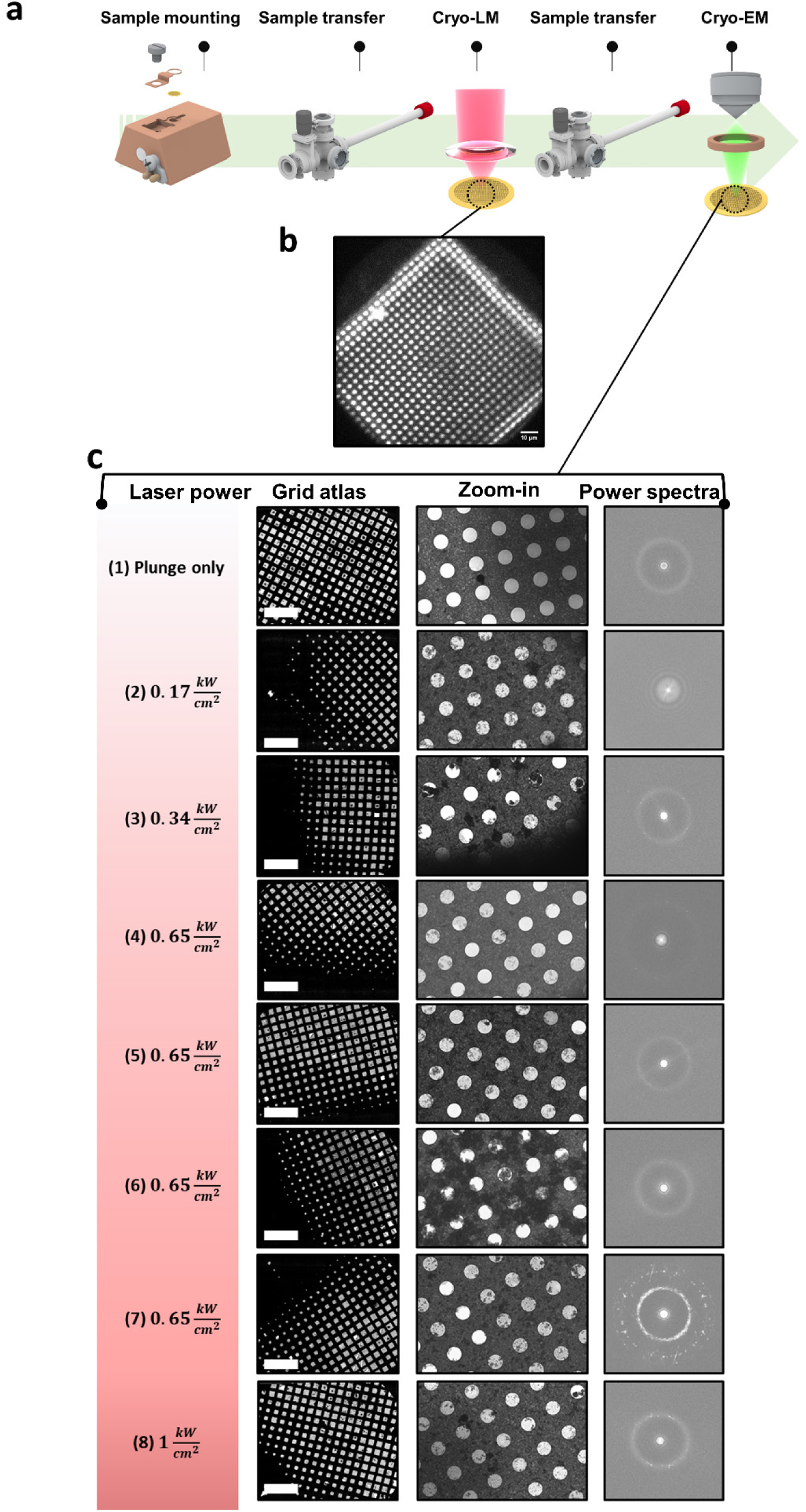
**Vitreous ice stability**. **a**, Pipeline of measuring vitreous ice stability after laser illumination. **b**, An example of fluorescence image of a single grid square recorded in the cryostat. The bright regions are holes containing frozen aqueous solution with nM fluorescent molecules. Samples are then retracted in reverse order to that shown in Fig. 1 and taken to a cryo-electron microscope (Titan Krios G2, 300 kV, K3 EFTEM) for investigation. **c**, Cryo-EM analysis of TEM grids exposed to different laser intensities, as noted on the left-hand side of each row (1–8). Micrographs of each TEM grid pattern (left column), the holey gold foil at intermediate magnification (middle column), and the associated power spectra (right column) of vitreous ice exposures in holes at high magnification (105k×, equivalent to 0.85 Å/px). Scale bar of the left column is 500 μm, and the hole size in the middle column is 2 μm. Some crystalline ice formation could be observed at various locations on most grids, independently of the laser exposure state. This is most likely due to the presence of transfer ice that accumulated during different steps (dewar transfer, grid clipping process, microscope transfer) of sample preparation.

**Supplementary Fig. 7.**
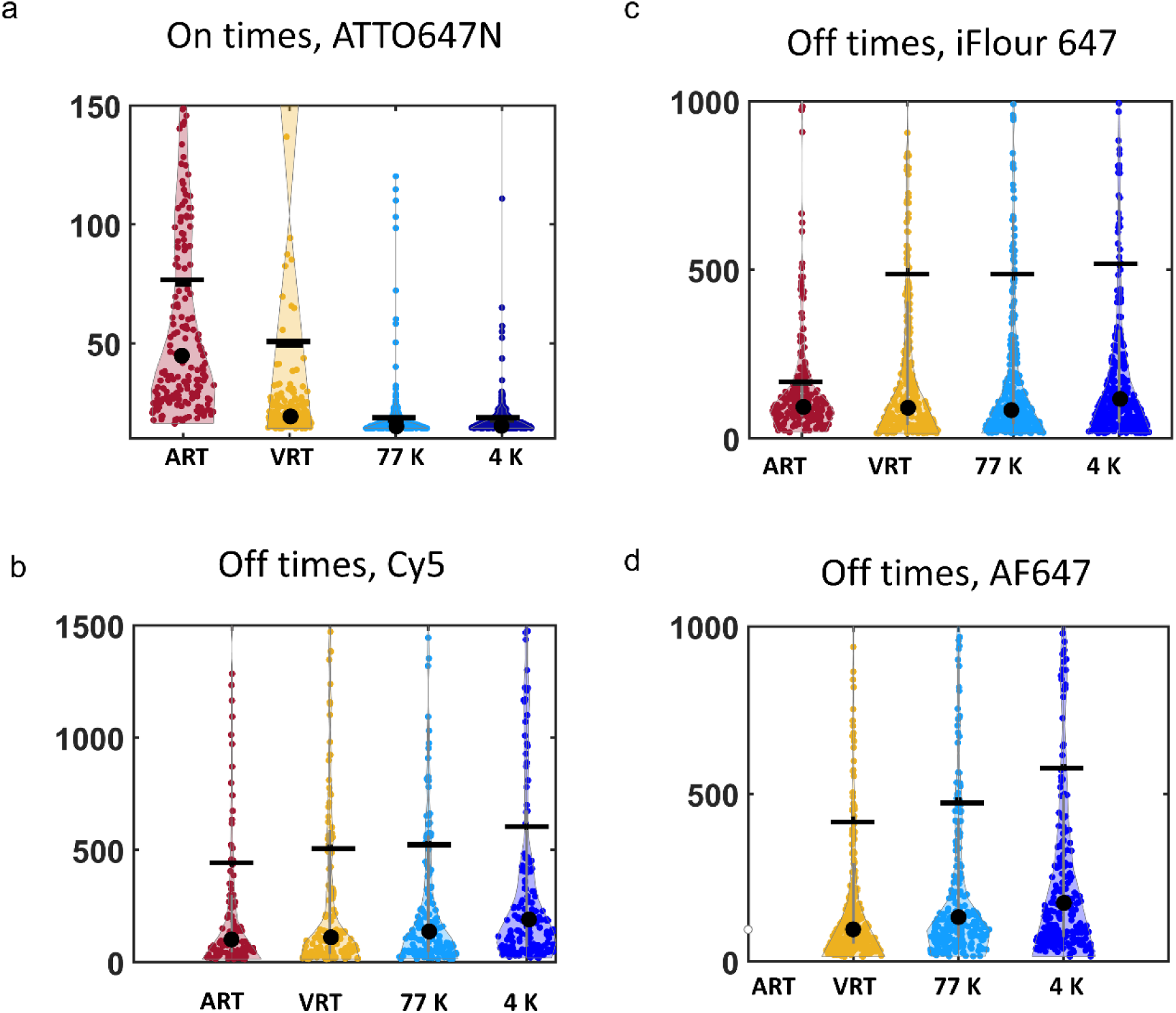
Photophysics characterization. **a**, Distribution of on dwell-times for ATTO647N recoded at different conditions (atmospheric room temperature (ART), vacuum RT (VRT), Vacuum 77 K, and Vacuum 4 K). **b-d**, Distribution of off dwell-times for different organic fluorophores: iFlour647 (b), Cy5 (c), Alexa Flour 647 (d). ART in (d) was not measured due to fast photobleaching of the sample. Black line indicates the mean values, and black circles indicate the medians.

**Supplementary Fig. 8.**
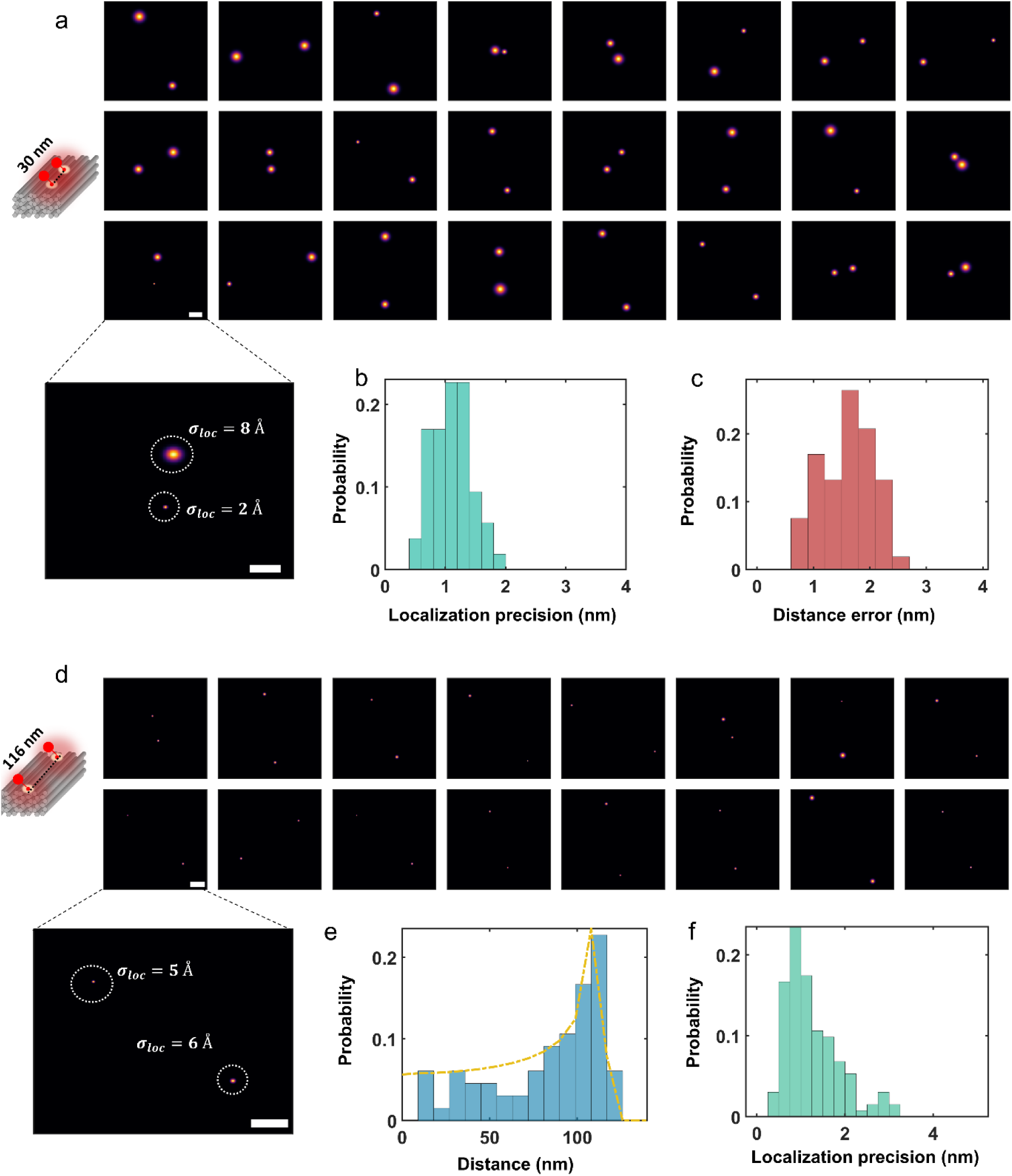
DNA nanoruler analysis and validation. **a**, Multiple 2D images of DNA nanoruler with two fluorophores separated by 30 nm**. b**, Localization precision histogram corresponding to measurements in (a)**. c**, Distance error estimated based on the localization precision described in Methods**. d**, Multiple 2D images of DNA nanoruler with two fluorophores separated by 116 nm**. e**, Pair-wise distance histogram (N=97) with a fit (yellow line) which takes the 3D orientations into account. The fit yields a distance of 117 nm close to the expected value of 116nm**. f**, Localization precision histogram for measurements in (d).

**Supplementary Fig. 9.**
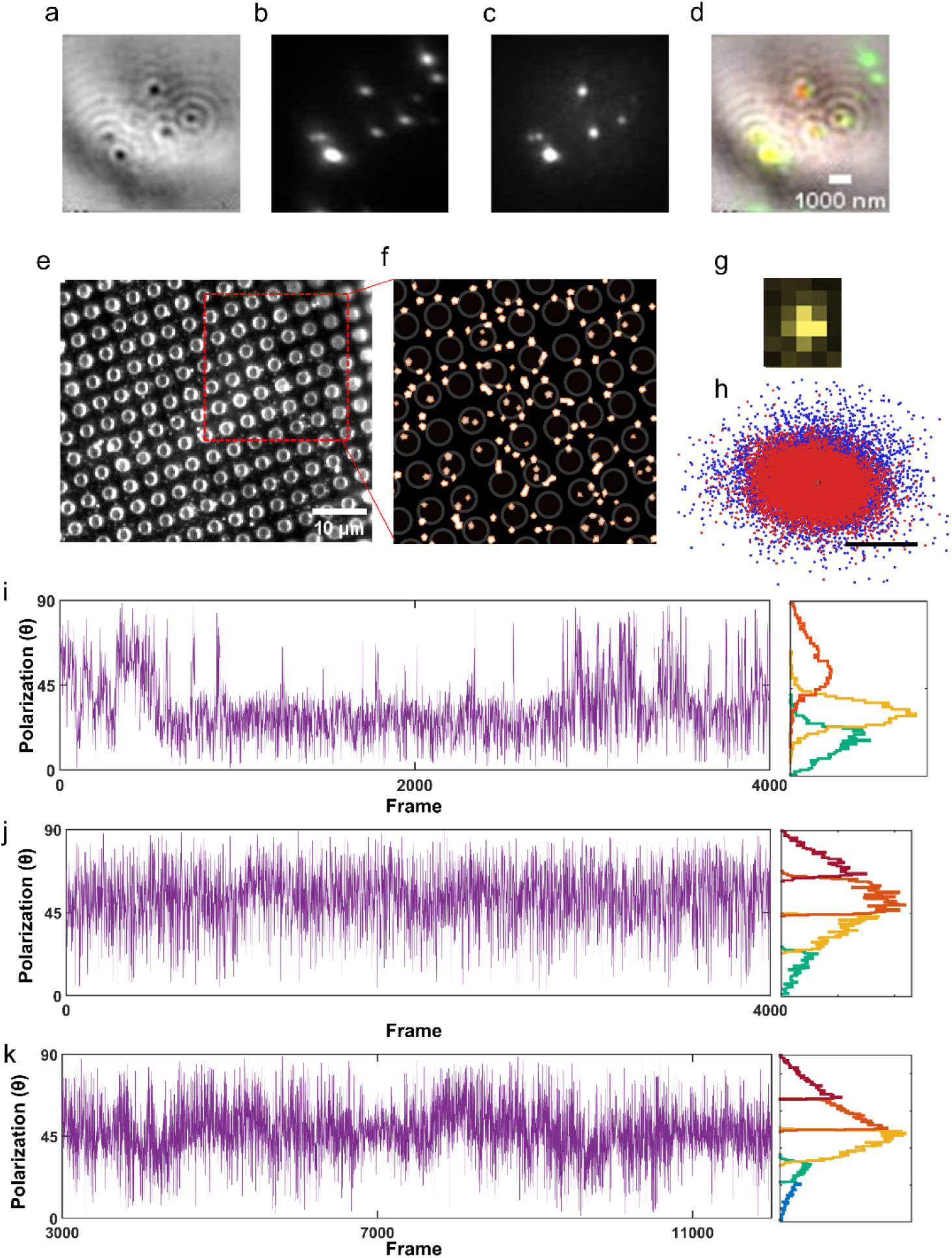
αHL characterization. To validate incorporation of αHL into lipid vesicles, we tethered the prepared sample (see Methods) onto a glass substrate functionalized with poly-Lysine. The samples were then imaged in a correlative fashion using interferometric scattering (iSCAT) microscopy and fluorescence microscopy. **a**, Signal from the iSCAT channel, clearly resolving the vesicles. **b**, To confirm that this signal indeed comes from vesicles, we labeled them with a fluorophore (memBrite 568). Spots coincide well with those in the iSCAT channel. **c**, Fluorescence image of αHL protein labeled with Alexa488-NHS ester. We can clearly see that the protein is localized within the vesicle signal. **d**, Registration of all channels further confirms the colocalization of the αHL signal (red color) with the vesicle signal (green color). **e-f**, TEM grid image of the protein in SLB recoded with our optical microscope shows sparse fluorescent molecules across the grid. **g**, An exemplary PSF. Each pixel is 227 nm. **h**, Overall localizations from the two polarization cameras (red and blue). **i-k**, Exemplary polarization time traces, showing 3-5 polarization components, as detected based on DSIC algorithm (*7*). We note that the signal is not continuous. It represents the on-times and discards the off-times.

**Supplementary Fig. 10.**
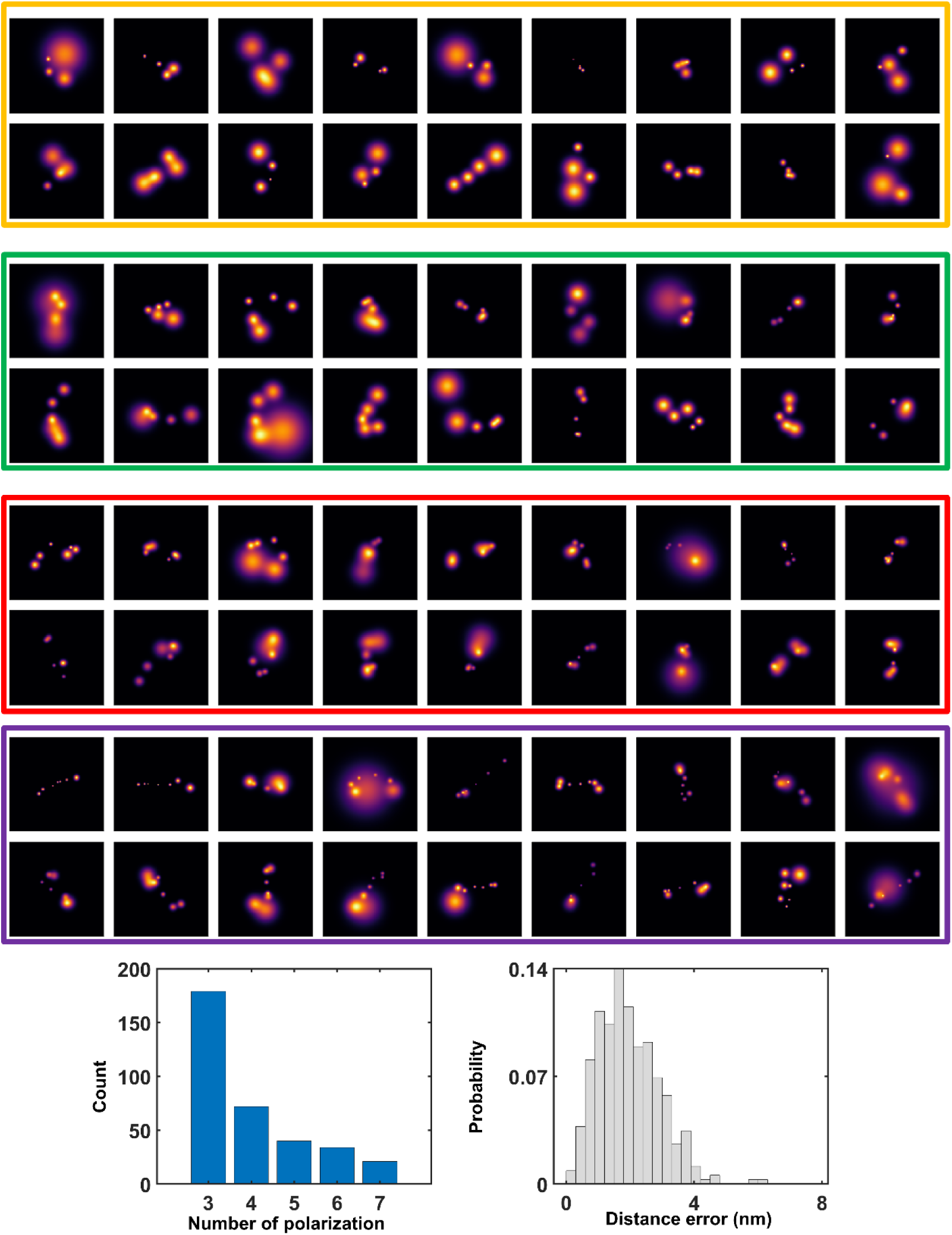
2D images obtained from an αHL sample. **a**, Some examples of the super-resolved 2D maps obtained from the polarization trace fitted with a model based on four-seven states. Particles with the orange, green, red and purples boxes show maps with 4, 5, 6, 7 fluorophore localizations, respectively. The particles show different projections in the sample plane with some restriction (see Figure 3). The image size is 120 × 120 pixel at 0.15 nm/pixel. **b**, Overall distribution of particles with more than two polarizations. **c**, Distribution of distance errors from the projections in (a).

**Supplementary Fig. 11.**
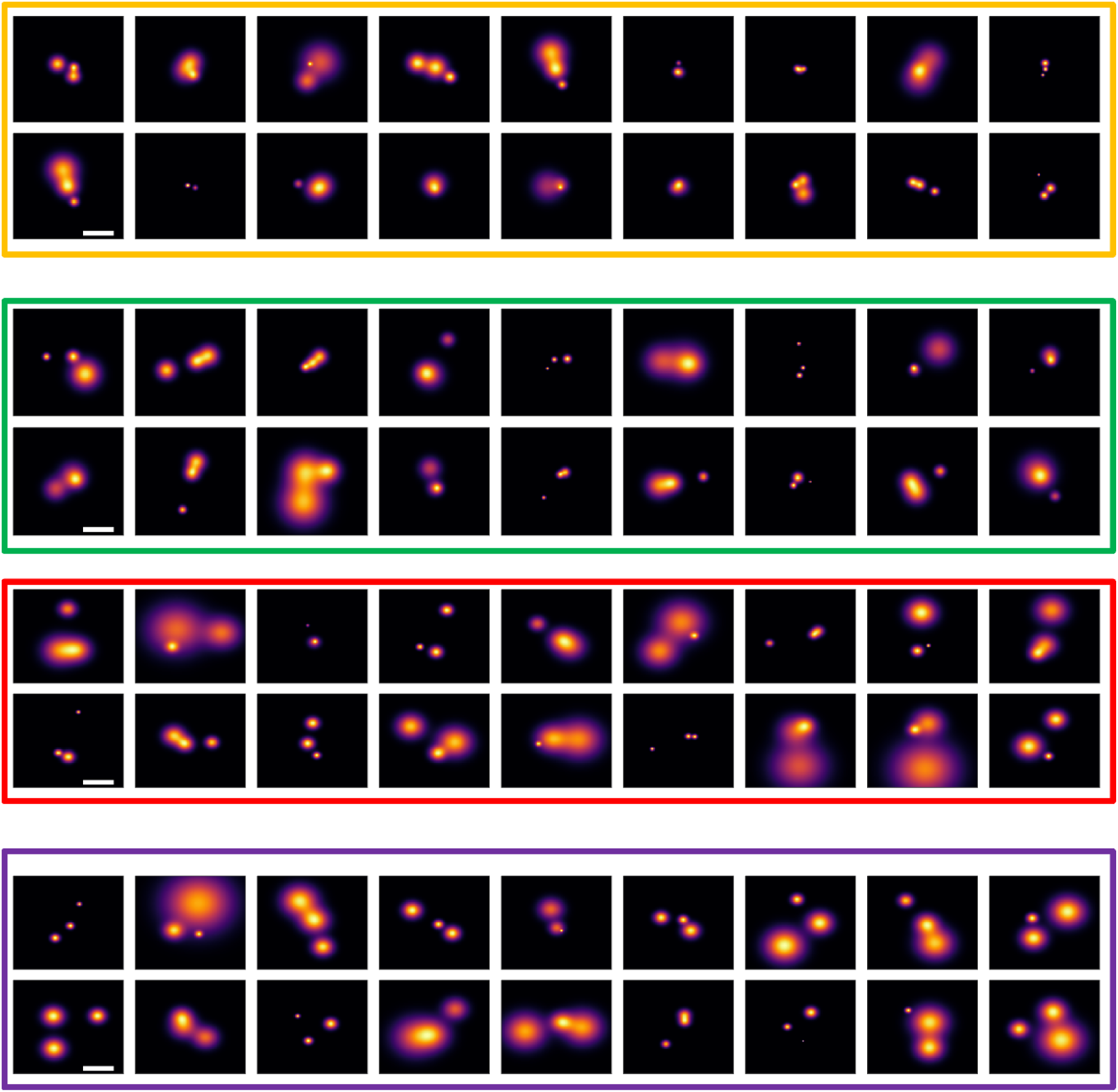
2D images of three fluorophores obtained from an αHL sample. Some examples of the 2D super-resolved maps obtained from polarization fitted with a model based on four-seven states. Particles were filtered based on localization precision below 3 nm and classified to the four inherent configurations as explained in Figure 3 of the main text (orange = class 1, green = class 2, red= class 3, purple = class 4). The image size is 120 × 120 pixel at 0.15 nm/pixel, and scale bar is 5 nm.

**Supplementary Fig. 12.**
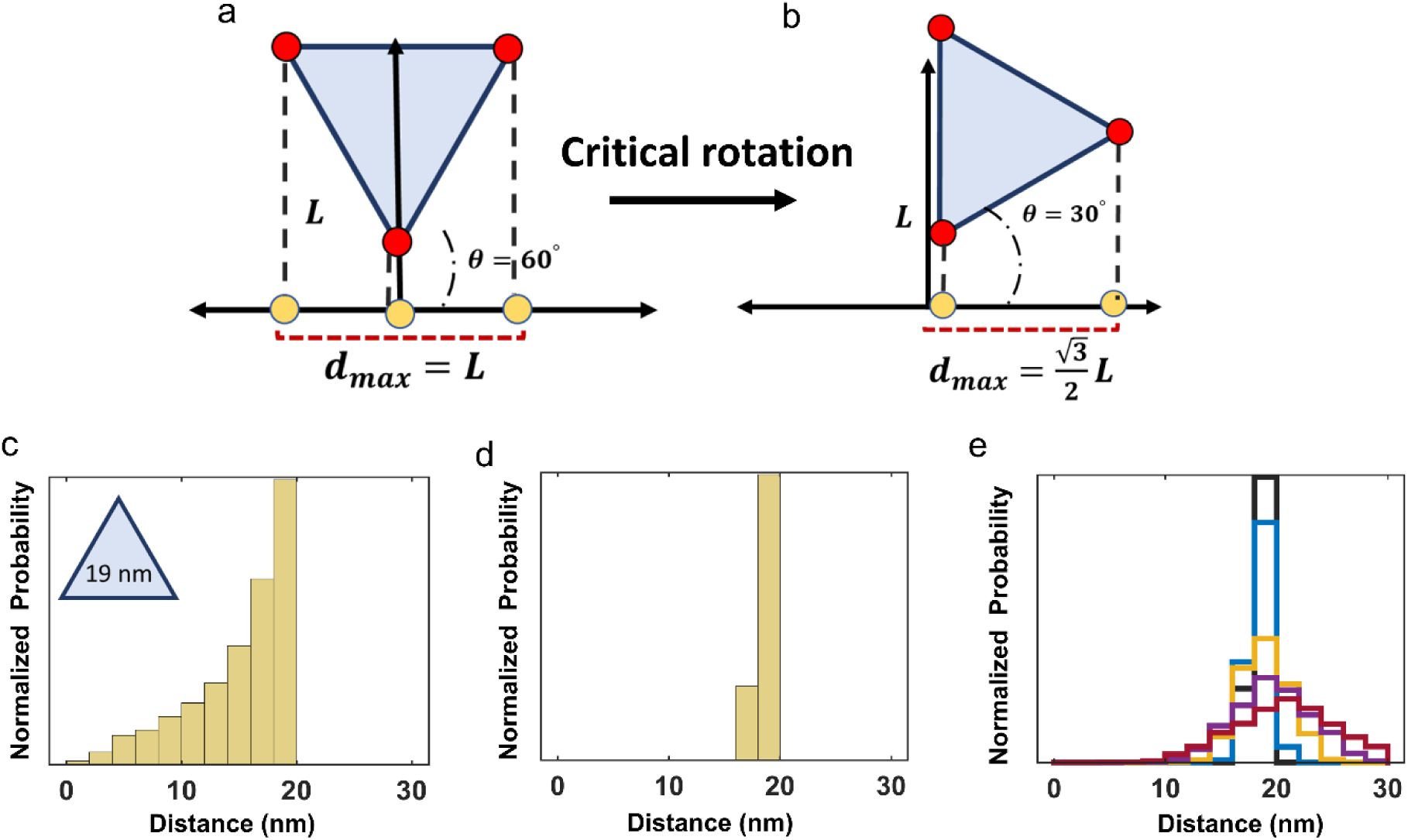
Pair-wise distance histogram of simulated data for an equilateral triangle versus localization precision. In general, if we consider a single distance *L* between two points, one could image the absolute distance if it lies in the image plane, namely at an angle (θ) of 0°. However, in case of tilting the sample, the projected distance onto the image plane becomes skewed toward shorter values, making the distance fall within a range from [0, *L*]. For three points in space forming a triangle (e.g. equilateral triangle), at least one of the projected side-lengths remains close to its true value regardless of the orientation. **a**, Scenario of an equilateral triangle with side length L. Projection onto the 2D plane yields side-length of L, and L/2. **b**, Scenario of another special orientation. projection yields side-lengths of 0 and 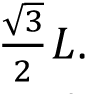 **c**, Pair-wise distance histogram generated from many randomly oriented triangles with side-lengths of 19 nm **d**, If we consider the maximum side length only from each rotated triangle, one obtains a value that is close to the true distance, resulting in a narrow distance distribution. The minimum distance in such a case is achieved at a critical angle of 30°, where the maximum distance reaches 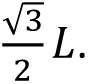 Distance histogram after considering the maximum side-length only. The histogram is notably narrow at the right side-length. In both cases in (c-d), we assumed perfect localization precision, 0. The width of such a histogram is strongly influenced by the localization precision of the data. The larger the localization precision, the broader is the peak. **e**, Histogram of maximum side-length as a function of localization precision. The decrease in the localization uncertainty from 0 to 0.7, 1.4, and 3 nm (color code: black, blue, orange, purple, and dark red, respectively) broadens the standard deviation of the distance distribution from 0.4 to 1.6, 3, and 5 nm, respectively. Therefore, in order to obtain valuable information on the protein scale, one must reach a precision that is superior to ∼ 2 nm.

**Supplementary Fig. 13.**
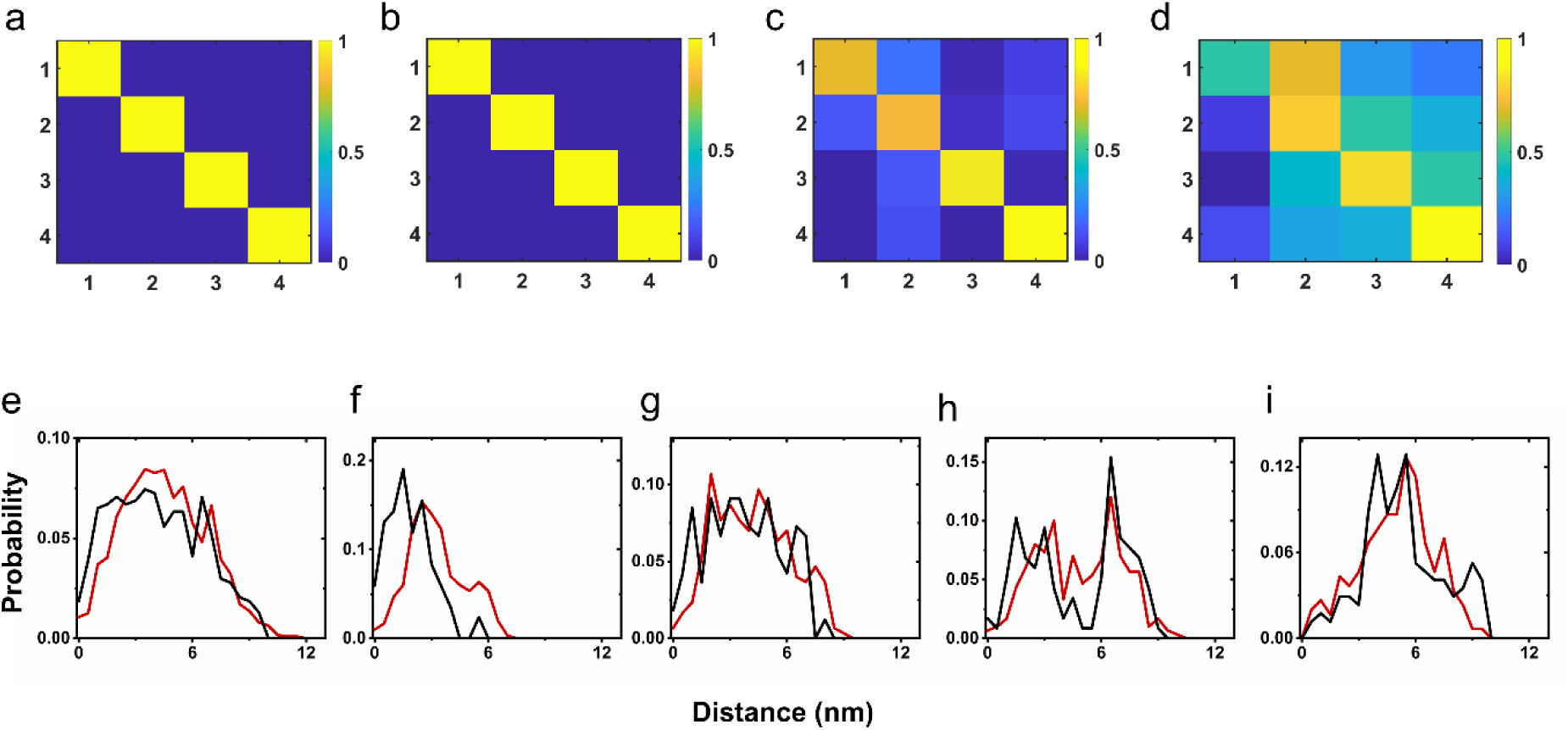
Particle classification and distance histogram fit of the αHL protein. Particle classification scores for the four inherent classes of αHL (as described in Fig. 4 of the main text). **a**, Classification using perfect localization (0 nm precision), showing accurate classification for all classes. **b**, Classification with 0.7 nm localization precision, maintaining accurate class assignments. **c**, Classification with 1.8 nm localization precision, where only classes 3 and 4 maintain perfect classification. **d**, Scenario with ±1 nm center displacement and 0.7 nm localization precision, showing degraded classification accuracy for classes 1 and 2. **e-i**, Distance histograms calculated from experimental data (black curves) for particles with three fluorophores, fitted with a model accounting for particle orientation and localization uncertainty (red curves; see Methods section in main text). (e) Overall distance histogram from unclassified particles, (f) Class 1 only, (g) Class 2 only, (h) Class 3 only, (i) Class 4 only.

**Supplementary Fig. 14.**
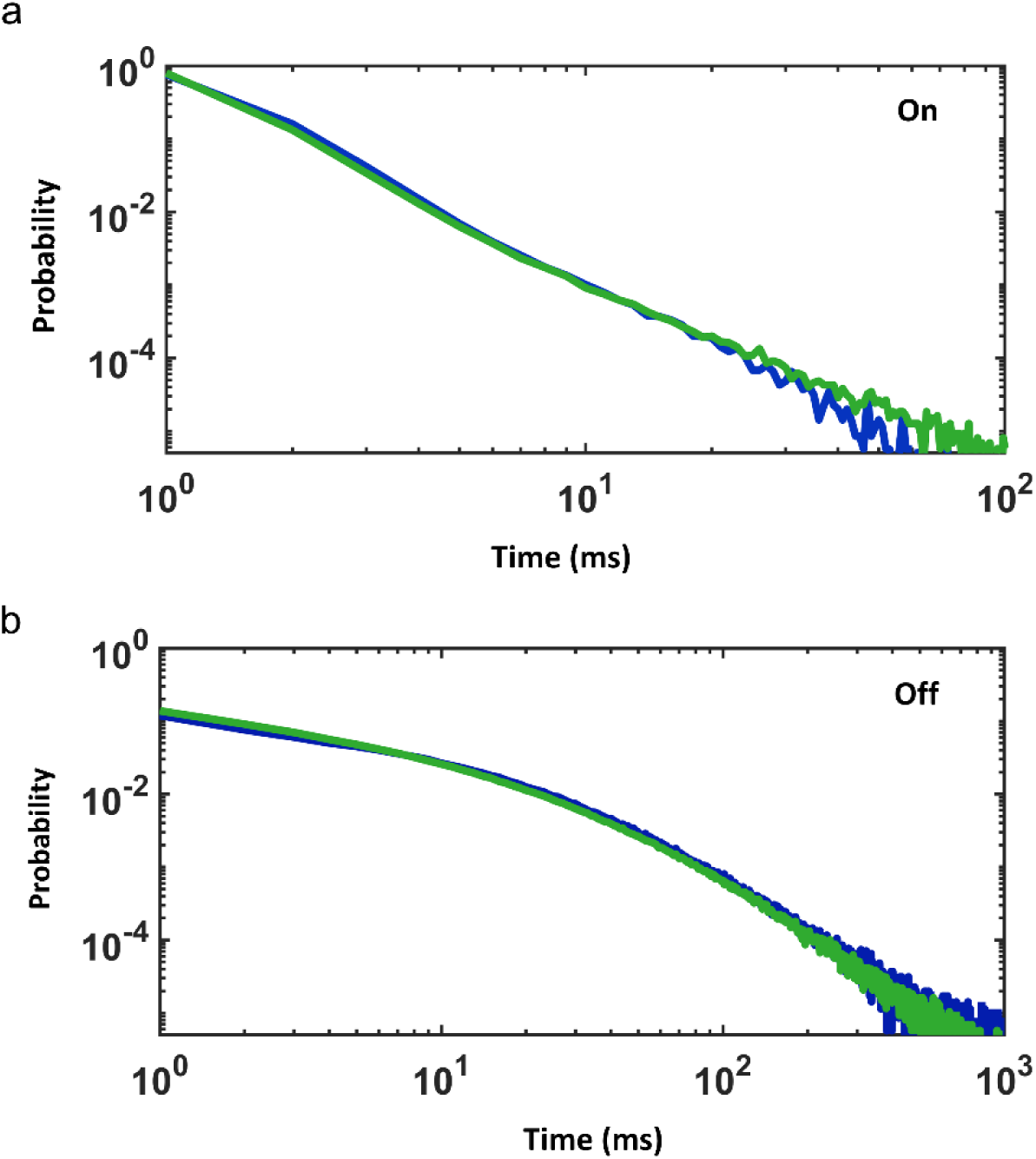
Comparative Photophysical Analysis of ATTO647N in Polyvinyl Alcohol (PVA) versus Vitreous Ice Matrices. **a**, On-times distribution of ATTO647N in PVA (blue curve) compared to that in vitreous ice (green curve) at high vacuum and LHe temperature. B, Off-times distribution. Color code and condition is the same as indicated in (a).

## Supplementary protocol

### Procedure at a Glance

| Step | Description |
| --- | --- |
| 1 | Evacuate the cryogenic optical microscope to a primary vacuum $<10^{-5}$ mbar. Start the flow of liquid helium into the cryostat until reaching 4 K. The vacuum at this stage goes down further to $<10^{-6}$ mbar. At this point, the optical microscope is ready for sample transfer. Keep the vacuum pump on and stabilize the helium flow to maintain a constant temperature. |
| 2 | Prepare the plunge-frozen TEM grids. Put them into storage in liquid nitrogen in a sealed grid box. |
| 3 | Purge the preparation chamber and the high-vacuum shuttle with dry nitrogen for ~10-20 minutes to minimize moisture. |
| 4 | Connect the transfer shuttle to the preparation chamber with its gate valve closed. Evacuate the transfer shuttle to a primary vacuum $<10^{-5}$ mbar and fill the dewar with liquid nitrogen. Wait for the temperature to stabilize at 90 K. From this point on, keep filling the dewar along all the procedure until the transfer is finished. (~ 15 min) |
| 5 | Prepare the preparation chamber. Move the glass vessel to the upper position and insert a cartridge into the cold stage adapter. Fill the vessel with liquid nitrogen until the cartridge is completely immersed and boiling stops. (~ 5- 10 min) |
| 6 | Quickly move the grid box from the storage container to the vessel. Insert a tweezer and a screw driver and wait for their tips to be cold. Maintain a constant flow of dry nitrogen gas through the chamber. (~ 1 min) |
| 7 | Carefully remove a grid from the grid box with the tweezer and place onto the recess in the cartridge. Fix the grid with the pre-cooled clamp by tightening the screw. (~ 1- 5 min) |
| 8 | Close the gate valve of the transfer shuttle and break the vacuum near the gate valve of the preparation chamber. Allow dry nitrogen to fill the atmosphere and the pipe of the vacuum pump to reduce the moisture. (~ 3 - 5 min) |
| 9 | Place the anti-contaminator plate on top the sample cartridge. Close the top part of the sample loading chamber and lower the glass vessel. Stop the dry nitrogen flow and start evacuating to a primary vacuum of $<10^{-1}$ mbar, open the gate valve of the shuttle and retrieve the cartridge with the manipulator. (~ 5 min) |
| 10 | Close the gate valve of the preparation chamber and switch on the turbo-molecular pump. Evacuate to a secondary vacuum of $<10^{-6}$ mbar. (~ 3 - 4 min) |
| 11 | Wait for the temperature and pressure to stabilize, then close the gate valve and switch off the pump. The vacuum in the shuttle is now maintained by the cold stage. |
| 12 | Connect the shuttle to the interface on the microscope (KF50 flange) and switch on the pump. Wait for the pressure in the differential pumping stage to reach $<10^{-5}$ mbar before opening the gate valves. (~ 2 min). |
| 13 | Open the gate valve of the shuttle first and let the pressure stabilize. Then open the gate valve of the microscope. (~ 2 min) |
| 14 | Insert the cartridge into the microscope cold stage adapter with the linear manipulator. Disconnect by rotating the manipulator to remove the screw from the thread in the cartridge. (~ 1 min) |
| 15 | Retract the manipulator, close the gate valves, vent the differential pumping stage and disconnect the shuttle from the microscope. Also disconnect the vacuum pump from the optical microscope. |
| 16 | Wait for at least 1-2 h before starting image acquisition. |

## Notes

### Competing Interest Statement

The authors have declared no competing interest.

### Funding

- Max Planck Society

